# Endocytome profiling uncovers cell-surface protein dynamics underlying neuronal connectivity

**DOI:** 10.1101/2025.08.28.672852

**Authors:** Colleen N. McLaughlin, Hui Ji, Katherine X. Dong, Chuanyun Xu, Kenneth Kin Lam Wong, Zhuoran Li, David J. Luginbuhl, Charles Xu, Cheng Lyu, Wei Qin, Jiefu Li, Namrata D. Udeshi, Steven A. Carr, Alice Y. Ting, Liqun Luo

## Abstract

Endocytosis actively remodels the neuronal surface proteome to drive diverse cellular processes, yet its global extent and circuit-level consequences have defied comprehensive interrogation. Here, we introduce endocytome profiling: a systematic, cell-type-specific approach for mapping cell-surface protein (CSP) dynamics in situ. Quantitative proteomic analysis of developing olfactory receptor neuron (ORN) axons generated an endocytic atlas comprising over 1,100 proteins and revealed the extent to which the surface proteome is remodeled to meet distinct developmental demands. Targeted interrogation of a junctional CSP showed that its endosome-to-surface ratio is precisely balanced to enable developmental axon pruning while preserving mature axon integrity. Multi-omic integration uncovered wide-spread transcellular signaling and identified a growth factor secreted by neighboring neurons to direct ORN axon targeting via endocytic regulation of its receptor. Endocytome profiling thus provides unprecedented access to cell-surface proteome dynamics and offers a powerful platform for dissecting proteome remodeling across diverse cell types and contexts.

---

**Neurons** must rapidly adapt to evolving developmental signals and physiological needs. The plasma membrane is the interface between neurons and their environment. Here, cell-surface proteins (CSPs) transduce external cues into the intracellular responses that control neurodevelopment and physiology. Transcriptional changes^1–5^ govern extensive, long-timescale shifts in the CSP landscape observed between developing and mature neurons^6,7^. However, neurodevelopmental processes, such as neurite growth and retraction, occur on much shorter timescales and thus require more rapid regulatory mechanisms. Targeted, on demand surface proteome remodeling is well-poised to shape these events. Although proteome-scale dynamics are increasingly appreciated in human cell lines^8–10^, the degree and impact of rapid cell surface remodeling in neurons or any cell *in vivo* is unknown.

Endocytosis actively sculpts the cell surface proteome. Within minutes^11^, soluble and membrane-anchored proteins are internalized from the plasma membrane into intracellular vesicles which facilitate their degradation or redistribution throughout the cell. These trafficking events modulate CSP signaling and adhesive functions, and depending on the context, can lead to signal attenuation, reactivation, or engagement of new molecular partners^12,13^. Mechanistic studies have elucidated the post-endocytic itineraries for select CSPs and revealed essential roles in processes ranging from neuronal survival^14^ to axon guidance^15,16^ and neuronal polarity^17^. However, these examples represent only a small fraction of the neuronal surface proteome. Thus, the extent of CSP endocytosis and identity of internalized cargos remain unknown for most cell types and developmental contexts. This gap stems, in part, from a lack of tools for simultaneous, quantitative comparison of both plasma membrane and endosomal compartments within the same cell type and physiological state.

Here, we present endocytome profiling, a proximity labeling approach that captures the dynamic protein exchange between the plasma membrane and endosomes within genetically defined cell types in the brain. We applied this method to profile a sub-population of ax-ons during development, a stage that requires reorganization of CSP-mediated adhesion and signaling. Quantitative proteomic analysis revealed that parallel profiling of the surface and endosomes is essential to determine each protein’s primary residence and yielded a compartment-resolved proteomic map comprising endolysosomal residents, CSPs, and secreted proteins. We leveraged this data to uncover a novel mechanism by which endocytosis remodels the CSP landscape to control the refinement and long-term stability of neural circuits. Finally, we demonstrate that this dataset can be used to identify transcellular signaling events that shape circuit connectivity. Endocytome profiling thus provides a powerful tool for decoding proteome-scale dynamics across cell states and systems.

## RESULTS

### Endocytosis is critical for target selection of olfactory receptor neurons

CSP endocytosis is known to influence axon guidance^15,16^; however, its role in the subsequent step of target selection—where neurons choose specific postsynaptic partners from many—remains unclear. We investigated this using *Drosophila* olfactory receptor neurons (ORNs), the primary sensory neurons of the olfactory system. Upon entering the brain, axons of the ∼50 molecularly distinct ORN subtypes follow a specific trajectory and target the dendrites of their postsynaptic partners, one of ∼50 subtypes of projection neurons (PNs) to form one-to-one synaptic connections within discrete and anatomically stereotyped units called glomeruli (Figure 1A).

**Figure 1.**
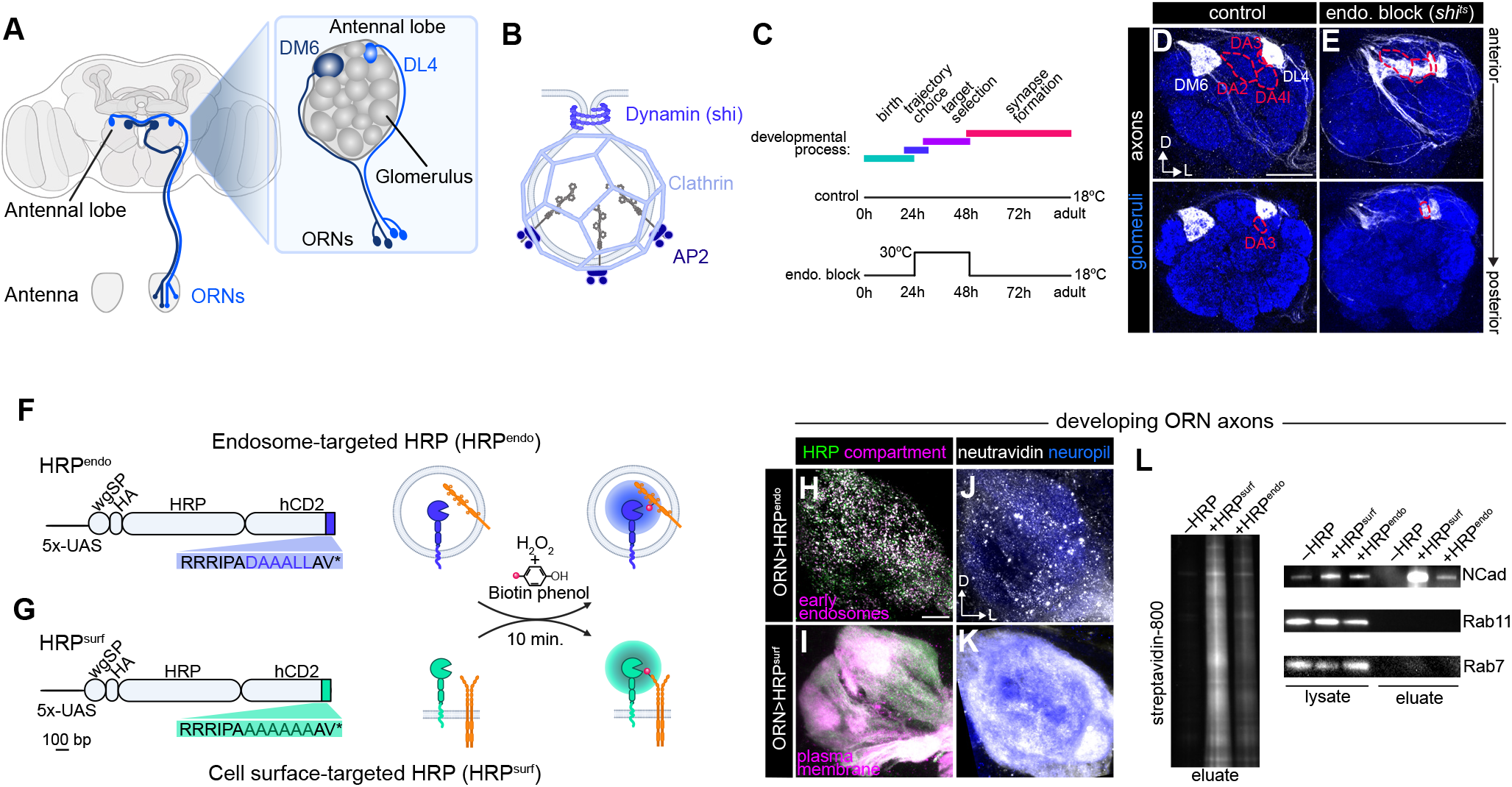
Endocytic control of olfactory receptor neuron axon targeting and development of endocytome profiling tools. (**A**) Adult *Drosophila* brain schematic highlighting the olfactory system (left) and the antennal lobe depicting the axon trajectories of two ORN subtypes and their terminals in the DM6 and DL4 glomeruli (magnified on the right). (**B**) Schematic of an endocytic vesicle and the molecular machinery necessary for clathrin-mediated endocytosis. (**C**) Schematic of ORN development (top) and the temperature shift paradigm used in controls (middle) or to inhibit dynamin function during axon target selection (bottom). Hours after puparium formation (h) are 25ºC equivalent. (**D, E**) Confocal images of ORN axons innervating the DM6 and DL4 glomeruli (white, labeled with membrane-tagged tdTomato) that express *UAS-shi*^*ts*^ via the *AM29-GAL4* driver of controls (D) and experimental flies (E). Dotted outlines denote glomeruli where ectopic targeting is observed. Scale bar, 20 µm. (**F, G**) Endosome-targeted (F) and plasma membrane-targeted (G) HRP transgenes (left) and labeling paradigm (right). wgSP, wingless signal peptide. bp, base pairs. Note that we use membrane-permeable biotin phenol to access intracellular compartments, instead of membrane-impermeable BXXP in our previous cell-surface proteomic profiling^6,7^. (**H**– **K**) Airyscan super-resolution images of 36h APF antennal lobes where all ORNs (via *Peb-GAL4)* express either *UAS-HA-HRP*^*endo*^ colocalized with early endosome marker *UAS-mCherry-2xFYVE* (H) or *UAS-HA-HRP*^*surf*^ colocalized with *UAS-myristoylated-RFP* (I). Neutravidin staining when *HRP*^*endo*^ (J) or *HRP*^*surf*^ (K) was expressed in ORN axons. Scale bar, 10 µm. (**L**) Post-enrichment bead eluate probed with streptavidin (left) and immunoblots of lysate and post-enrichment eluate stained for NCad and Rab proteins (right). In this and all subsequent Figures: D, dorsal; L, lateral. NCad (in blue) is used to label neuropil/glomeruli. * p < 0.05; ** p < 0.01; *** p < 0.001; **** p < 0.0001; ns, not significant. n, the number of the antennal lobes quantified. Images were from antennal lobes of young adults (< 5 days old), unless otherwise noted. Detailed information regarding all genotypes can be found in Table S1. See Figure S1, 2 for additional data.

Clathrin-mediated endocytosis is the primary receptor internalization pathway in neurons. It involves the sequential recruitment of adaptor proteins such as AP2 to cluster endocytic cargos, followed by clathrin and GTPase Dynamin which mediates vesicle scission from the plasma membrane^18,19^ (Figure 1B). Since earlier axon guidance decisions, such as trajectory choice, can influence target selection^20^, we temporally restricted endocytic inhibition to the latter process by expressing a temperature-sensitive *shibire* transgene (*shi*^*ts*^, encoding a dominant negative form of Dynamin) only in ORN axons projecting to the DM6 and DL4 glomeruli (Figure 1A, D). *shi*^*ts*^ only blocks Dynamin function when flies are at high temperatures (29–31ºC)^21^, giving us precise control of when endocytosis is disrupted.

Rearing these flies at 30ºC for a 24-hour window after the majority of axons have selected their trajectory^22^ but before target selection and synapse assembly^23^ resulted in robust and specific defects in ORN targeting (Figure 1C–E). DM6-ORN axons exhibited a significant reduction in targeting to their normal glomerulus and a significant increase in innervation to two specific glomeruli, DA2 and DA4l, when Dynamin activity was blocked (Figure 1E, S1A). Similarly, impairing expression or function of AP2 or clathrin led to stereotyped mistargeting of DM6-ORN axons (Figure S1C, D). Likewise, blocking dynamin activity in DL4-ORNs caused them mistarget to the DA3 glomerulus (Figure 1E, S1B). The fact that endocytic impairment caused ORN axons to innervate stereotyped, but incorrect, glomeruli, indicates that this process regulates a specific repertoire of CSPs necessary for axon target selection.

### Proximity labeling tools for profiling endocytic and cell-surface proteomes

To comprehensively define the CSPs enriched in endosomes during axon targeting, we developed a quantitative proximity labeling approach to chart the surface-to-endosome distribution of CSPs in developing ORN axons. CSP selection for clathrin-mediated endocytosis can occur via short amino acid motifs in their cytoplasmic tails^24,25^. Thus, we targeted the proximity labeling enzyme horseradish peroxidase (HRP) to the lumen of endosomes, hereafter HRP^endo^, by placing it N-terminal to the extracellular domain of human CD2 (a single-pass transmembrane protein) and adding a dileucine-based endocytic motif to its cytoplasmic tail (Figure 1F). The entire open reading frame is under *UAS* control for GAL4-mediated cell-type-specific expression (Figure 1F). We validated that HRP^endo^ was internalized *in vitro* and *in vivo* (Figure S2A, C), and that it colocalized with markers of the endolysosomal system (Figure S2B, D–F). Further, in developing ORN axons, HRP^endo^ partially colocalized with mCherry-2xFYVE, which predominantly accumulates on early endosomes^26^ (Figure IH), validating that it was in the correct subcellular compartment in our cells of interest.

We labeled CSPs residing on the plasma membrane by mutating the dileucine motif of HRP^endo^ to alanines, hereafter referred to as HRP^surf^ (Figure 1G). HRP^surf^ should take the same secretory route as HRP^endo^ to the plasma membrane but should not robustly enter endosomes via clathrin-mediated endocytosis. We validated that HRP^surf^ was predominantly at the plasma membrane in ORN axons and cultured cells (Figure 1I and S2A, B).

Upon H_2_O_2_ application, both HRP^endo^ and HRP^surf^ labeled proteins using biotin-phenol in dissected brains (Figure 1J, K). The biotin distribution reflected their respective subcellular compartments, with HRP^endo^ labeling appearing punctate and HRP^surf^ being continuous throughout ORN axons (Figure 1J, K). Streptavidin bead enrichment confirmed labeling of a wide-range of proteins (Figure 1L, left), and immunoblots validated compartment specificity—detecting the neuronal surface protein N-Cadherin (NCad), but not cytosolic endosome-associated Rab GTPases, Rab7 and Rab11 in streptavidin bead eluates (Figure 1L, right). These results are consistent with HRP’s orientation toward the extracellular and luminal space, and the membrane impermeability of the biotin-phenoxyl radical it generates, which prevents labeling of cytosol residing proteins, such as Rabs.

### The axonal endocytome and surfaceome are enriched for distinct proteins

Next, we combined HRP-mediated biotinylation with mass spectrometry-based proteomics to define the ‘endocytome’ and ‘surfaceome’ of ORN axons. We specifically profiled ORN axons during target selection, as we could easily dissect distal axons away from their cell bodies in the antenna (Figure 1A,C). Using a tandem mass tag (TMT)-based quantitative strategy^27^, we profiled three biological replicates of HRP^endo^ or HRP^surf^, alongside two negative controls (lacking either HRP or H_2_O_2_) to account for non-specific labeling (Figure 2A).

**Figure 2.**
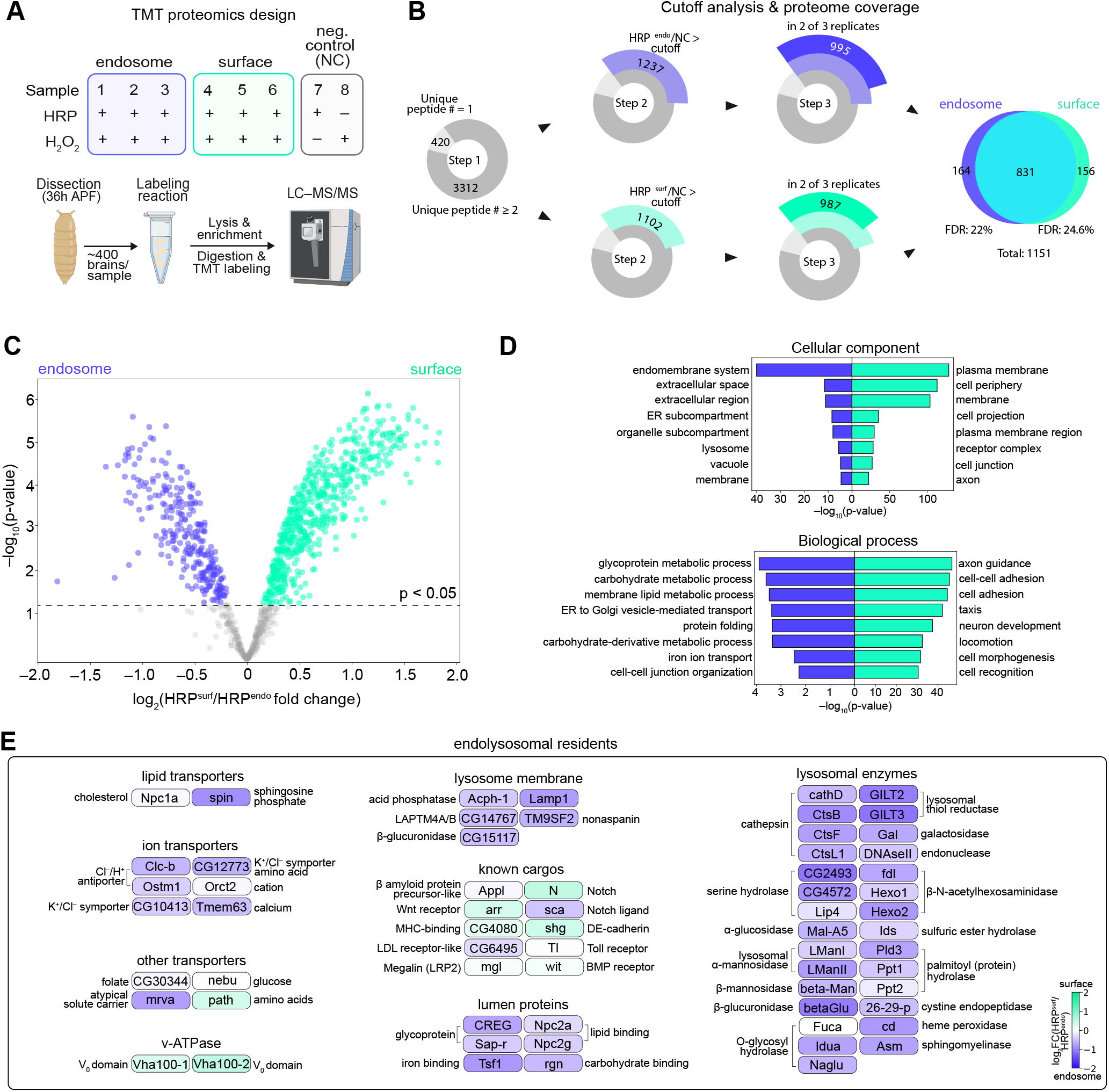
Proximity labeling enables profiling of global CSP dynamics in ORN axons. (**A**) Design of the tandem mass tag (TMT)-based quantitative proteomics experiment. Each genotype comprises three biological replicates (blue or green) in addition to two negative controls (gray). (**B**) Summary of cutoff analysis applied to the proteomes and number of proteins retained after each filter. FDR, false discovery rate. (**C**) Volcano plot depicting individual protein enrichment in the axonal endocytome or surfaceomes. Dashed line denotes p-value cutoff applied to proteomes yielding 608 surfaceome and 267 endocytome proteins used for Gene Ontology (GO) analysis. Moderated p-value is depicted as cutoff. Note: TMT-based proteomics can have ratio compression and result in an underestimation of true abundance differences between samples. (**D**) Top 8 GO categories cellular component (top) and biological process (bottom). (**E**) Schematic of proteins known to be in the endolysosomal system defined by Flybase (www.flybase.org), Park et al., 2022^91^ or Itzhak et al., 2016^8^. Color indicates level of enrichment endosome or surface enrichment (in log [HRP^surf^/HRP^endo^] fold change). We note that the surfaceome was enriched for three known endolysosomal residents including V_0_ vacuolar H^+^ -ATPase subunits and an amino acid transporter (path), both with known endolysosomal and plasma membrane localization^92,93^. See Figure S3 and Table S2 for additional data.

From the 3,732 total proteins detected by mass spectrometry, we identified compartment-enriched proteins via four filtering steps. First, we retained the 3,312 proteins with at least two unique peptides detected (Figure 2B; Table S2). Second, we rank-ordered proteins by their experimental-to-negative control TMT ratios in descending order and validated that proteins containing a signal peptide and/or a transmembrane domain were enriched, whereas putative contaminants lacking these signatures were not (Figure S3A–D). Following published protocols^6,28^, we filtered out contaminants using this ratiometric approach (Figure 2B). Third, since HRP^endo^ or HRP^surf^ replicates were highly similar (Figure S3E), we retained proteins found in at least 2 of 3 proteomes per condition (Figure 2B). These steps yielded a total of 1,151 proteins, most of which were detected by both HRP^endo^ and HRP^surf^ labeling (Figure 2B), highlighting the dynamic nature of protein localization. Finally, to determine compartment-specific enrichment, we compared the extent of biotinylation of these 1,151 proteins using their relative abundance in HRP^endo^- vs. HRP^surf^ -labeled proteomes. This yielded 267 and 608 proteins that were significantly enriched in the endocytome and surfaceome, respectively (Figure 2C).

Confirming the spatial specificity of our approach, gene ontology (GO) analysis classified endocytome proteins within the extracellular region, endomembrane systems, and lysosomes, whereas surfaceome proteins localized to the plasma membrane and cell periphery (Figure 2D). Consistent with the stage profiled, the top biological process GO terms in the surfaceome were related to axon development and cell adhesion (Figure 2D). The endocytome was enriched for metabolic and ion transport terms (Figure 2D), in line with the role of this compartment in macromolecule degradation. Deeper investigation of the proteins within these categories revealed lysosomal enzymes, integral membrane proteins, and membrane transporters (Figure 2E), establishing a comprehensive atlas of endolysosomal residents and further validating our profiling. We also examined known endosomal cargos and found several to be differentially distributed between the plasma membrane and endolysosomal compartments, in line with cell- and context-specific nature of CSP endocytosis (Figure 2E). Notably, cell-cell junction organization terms were highly enriched in the endocytome (Figure 2D), implying that internalization of this specific class of proteins is important for axon development (see Figures 4, 5).

Our endosomal profiling also captured crosstalk between the secretory pathway and endolysosomal network as evidenced by the presence of endoplasmic reticulum (ER)-related GO terms (Figure 2D). Protein exchange between these two systems is well-documented^29,30^ and proteins that enter the endolysosomal system via biosynthetic vesicles typically participate in degradative processes^31,32^ or are themselves targeted for degradation^33^. In fact, several of these proteins have been implicated in a recently identified ER-to-lysosome degradation pathway^34–38^ (Table S2), suggesting that this may occur in axons. These findings highlight that endocytome profiling detects proteins entering the endolysosomal system from both the plasma membrane and biosynthetic compartments and lay the groundwork for future investigation of their fates and functions in developing axons.

### Dual-compartment profiling is required to resolve protein localization

Prior studies have used cell-surface proximity labeling to define long-timescale changes in the steady-state protein composition across development^6,7^ and identified internalized CSPs by profiling immunoprecipitated neuronal endosomes^39^. Although useful for characterizing compartment-specific proteomes, we reasoned that profiling each compartment in isolation would not only provide less quantitative insight into a protein’s distribution across compartments but could skew datasets toward highly abundant proteins, rather than those enriched in each one.

We directly tested the extent to which protein abundance impacts labeling by examining the top 200 transmembrane (TM) proteins labeled by HRP^endo^ before comparing them to HRP^surf^ (Figure 3A). Indeed, 62% of these proteins were also among the top 200 TM proteins labeled by HRP^surf^ (Figure 3A). This overlap led us to ask how these proteins were distributed between the surface and endosomal compartments. In fact, many were more enriched at the cell surface (Figure 3B), likely reflecting high overall protein levels. To validate these findings, we analyzed the localization of four representative TM proteins in ORN axons and found that their spatial distribution mirrored the compartment assignments from our ratiometric analysis (Figure 3B–D). These data highlight the dynamic nature of protein localization and emphasize that determining a protein’s primary residence requires concurrent profiling of both compartments.

**Figure 3.**
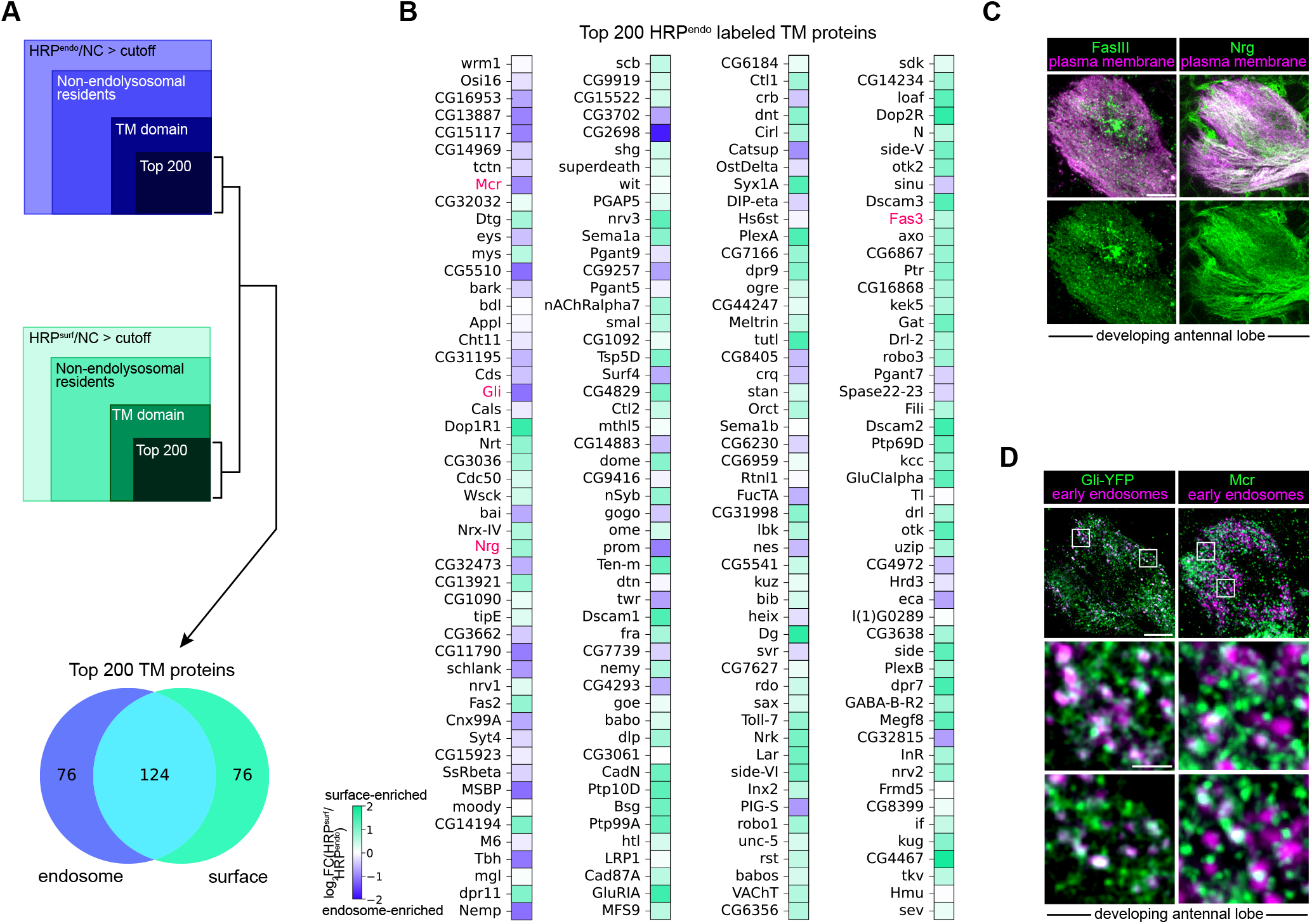
Dual-compartment profiling is required to resolve preferential protein localization. (**A**) Filters used to identify the top 200 transmembrane (TM) proteins labeled by either HRP^endo^ (top) or HRP^surf^ (middle) and a Venn diagram (bottom) of the proportion of the TM proteins that are labeled by either/both HRP transgenes. (**B**) Heatmaps depicting endosome and surface enrichment (in log_2_ [HRP^surf^/HRP^endo^] fold change) of the top 200 TM proteins labeled by HRP^endo^. Proteins are rank ordered from highest to lowest labeled by HRP^endo^ from top left to bottom right. Many of the top 200 TM proteins labeled by HRP^endo^ were more highly enriched in surfaceomes than endocytomes. Enrichment of TM proteins in pink was validated in C and D. Note: some of the top proteins are regulate secretory pathway-to-endolysosomal network trafficking or degradation (detailed in Table S2 and Figure 2). (**C**) Images of colocalization of myristoylated-RFP expressed only in ORNs and surface-enriched proteins FasIII (left) and Nrg (right). Note that based on anatomy and membrane staining, the puncta in FasIII images are largely from projection neurons. Images were taken at ∼36h APF. Scale bar, 5 µm. (**D**) Airyscan super-resolution images of all ORN axons expressing the early endosome marker mCherry-FYVE and endogenously tagged Gliotactin (Gli)-YFP (left) and anti-Mcr (right). Images were taken at ∼36h APF. Scale bar, 5 µm (D); 1µm (D, zoom). Note: non-colocalized puncta are likely in other endolysosomal compartments.

Collectively, these data establish proximity labeling as a powerful strategy for identifying proteins regulated by the endolysosomal network within a subset of axons in the intact brain. To showcase the utility of this tool in revealing how endocytic remodeling equips neurons to meet distinct developmental demands, we pursued two complementary approaches: (1) defining how the spatial distribution of CSPs instructs their function in developing and mature neurons; and (2) leveraging multi-omics to uncover cell-cell interactions underlying circuit assembly.

### The endocytome contains multiple cell-cell junction proteins required for axon development

We first focused on the CSP cargos most enriched in endosomes. In contrast to the surfaceome where many canonical neurodevelopment proteins were detected (Figure S4A, B), cell–cell junction CSPs were highly enriched in the endocytome (Figure 4A). These junctions include septate, tight, and adherens junctions and are formed by CSPs that link cells together or to the extracellular matrix^40^. In the nervous system, such junctions have been reported to form between progenitors, glia, and myelin and axons^41–43^, but not between neurons.

**Figure 4.**
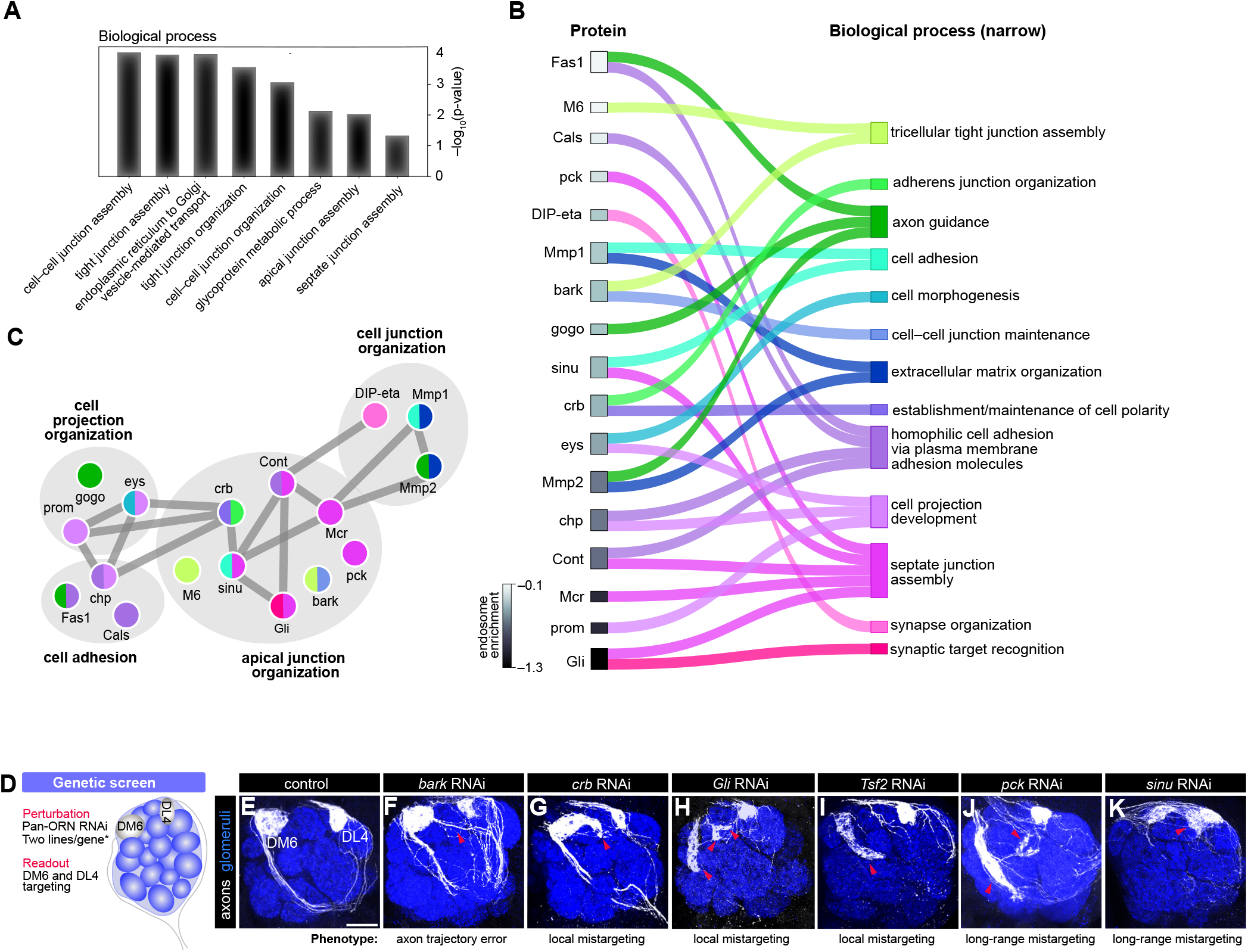
The endocytome contains multiple cell-cell junction proteins critical for axon targeting. (**A**) GO biological process analysis of endosome-enriched membrane proteins performed on proteins. (**B**) Chord plot depicting endosome-enriched proteins that fell within cell–cell junction categories in panel A and the specific process(es) they are associated with (right). Proteins are rank ordered from lowest to highest (top to bottom) endosomal enrichment. Gray scale in rectangles indicates level of enrichment (log_2_ [HRP^surf^/HRP^endo^] fold change). (**C**) STRING plot depicting cell–cell junction proteins as well as the other endosome-enriched CSPs they interact with. Links between nodes denote protein-protein interactions and node colors depict the category assignment(s) in B. (**D**) Schematic depicting loss-of-function screen genetic screen where membrane proteins enriched in the endocytome were knocked-down in all ORNs (via *peb-GAL4)* and axons projecting to the DM6 and DL4 glomeruli were labeled with *AM29QF2* driven membrane-targeted tdTomato (*QUAS-mtdTomato*; white). Scale bar is 20 µm. (**E**–**K**) Representative images of antennal lobes depicting phenotypes resulting from knockdown of each cell-cell junction protein in ORNs. The DM6-ORN axon phenotypes and their penetrance are detailed in Table S3. Arrowheads indicate DM6ORN axon mistargeting. See Figure S4 and Table S3 for additional data.

Since ORN axons have minimal glial contact during target selection^44,45^, the endosomal enrichment of these junctional proteins was unexpected.

This prompted us to perform focused analysis of proteins in the cell-cell junction, cell adhesion and cell projection organization categories, as many of them interact (Figure 4B,C). We annotated each protein with the specific biological process(es) it was most implicated in. A few proteins had known neuronal roles, but the majority did not (Figure 4B). Our prior colocalization experiments confirmed endosomal enrichment of two junctional proteins, Macroglobulin complement-related (Mcr) and Gli (Figure 3D).

Next, we conducted a targeted loss-of-function screen to evaluate if these junctional proteins could have functions in axon development. We expressed RNAi lines against each gene in all ORNs and monitored axons projecting to the DM6 glomerulus (Figure 4D). All six genes tested produced axonal phenotypes (Figure 4E–K; Table S3). For example, the scavenger receptor bark beetle (bark) promoted axon trajectory choice but did not affect later developmental events (Figure 4F). Loss of the adherens junction protein *crumbs* (*crb*) or septate junction components *Transferrin 2* (*Tsf2*) and *Gli* caused axons to mistarget to glomeruli nearby DM6 (local mistargeting defects; Figure 4G–I), whereas knockdown of the claudins *pickel* (*pck*) and *sinuous* (*sinu*) caused long-range mistargeting defects (Figure 4J,K). These data indicate that junctional CSPs control distinct aspects of axon targeting. Interestingly, a recent study found transcripts for tight and adherens junction CSPs enriched in mammalian neurons during neurite outgrowth and synaptogenesis^4^, suggesting that this class of proteins may act as conserved regulators of circuit assembly.

Since this is the first profiling of the ORN surfaceome, we also examined if surface-enriched CSPs regulate axon development. Five of the six genes we tested controlled one or more aspects of axon targeting (Figure S4C–I; Table S3). The fact that 91% (10/11) of the CSPs tested produced phenotypes validates both the biological relevance of our profiling and the idea that circuit assembly depends on a diverse set of CSPs distributed across endosomes and the plasma membrane.

### Crumbs promotes axon pruning and is endocytosed to safeguard axon integrity

How does endocytosis shape the neuronal roles of cell-cell junction proteins? We addressed this question by studying crb, an evolutionarily conserved protein involved in epithelial polarity^46,47^, adherens junction formation^48^, and photoreceptor integrity^49,50^. Humans have three CRB proteins, with CRB1 and CRB2 being most similar in size and domain composition to the single fly crb (Figure 5A,top). Though CRB1/2 are expressed in the brain^51,52^, no role for any crb ortholog has been described in this context.

**Figure 5.**
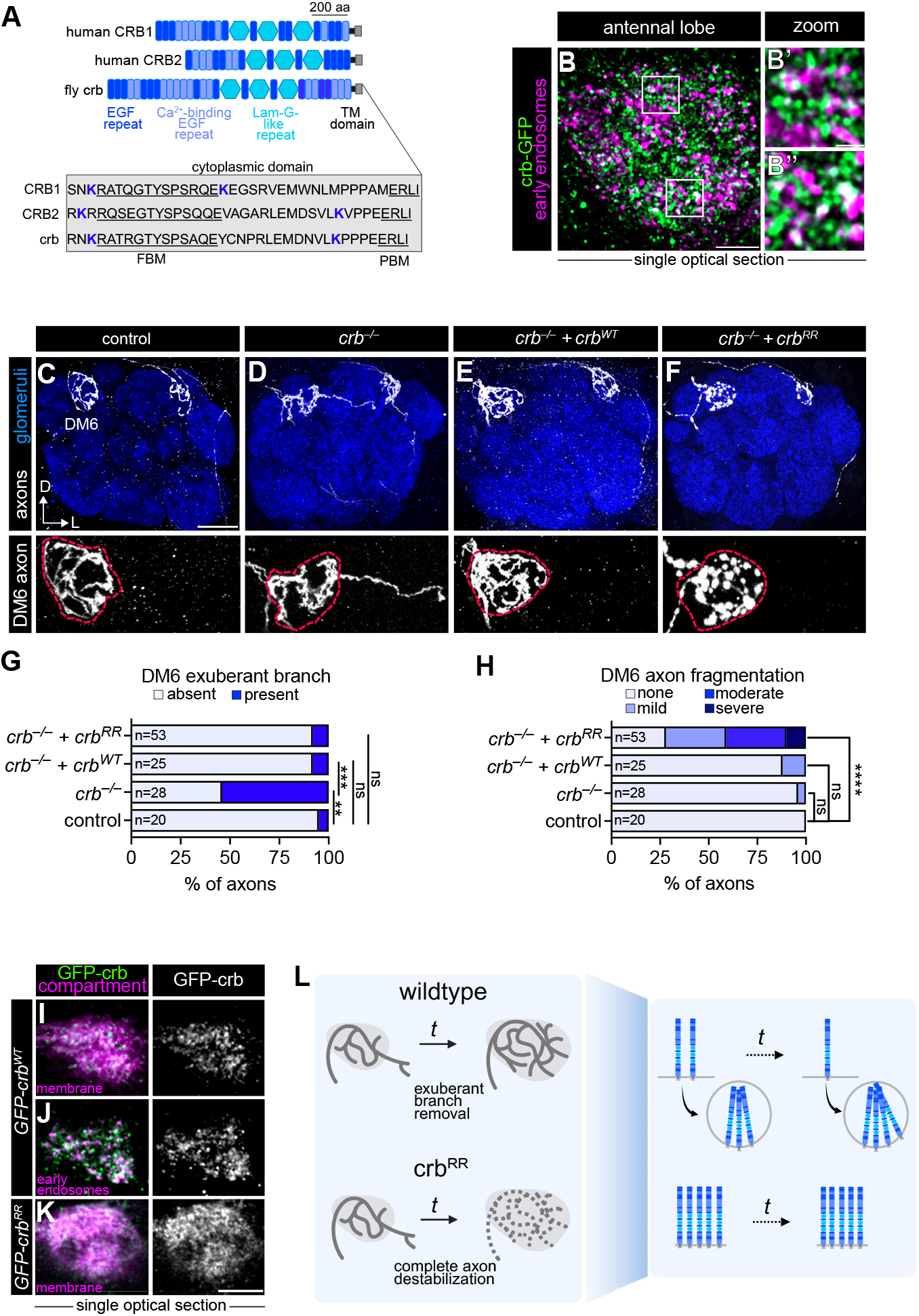
Crumbs promotes axon pruning and is endocytosed to safeguard axon integrity. (**A**) Top, domain structure (Uniprot) of human and *Drosophila* crumbs proteins. Bottom, the dcytoplasmic domain is highlighted in the gray box with the two lysines (K) that facilitate fly crb endocytosis in blue. FBM is FERM-domain binding motif, PBM is PDZ-domain binding motif. aa, amino acids. (**B**) Airyscan super-resolution image of all ORN axons expressing early endosome marker mCherry-FYVE and endogenously tagged crb-GFP allele. Zoomed-in images (B’, B’’) show crb-GFP colocalization with mCherry-FYVE. Images taken at ∼36h APF. (**C**–**F**) Images of single DM6 and DL4 axons of indicated genotypes from *hs-FLP-*based MARCM clones (top) and zoomed in images of the DM6 axons depicting membrane integrity (bottom). Dashed lines denote the DM6 glomerular boundary. (**G, H**) Quantification of the percentage of DM6 axons that still have the exuberant branch present at the adult stage (G) and axon fragmentation (H) of indicated genotypes. (**I**–**K**) Airyscan super-resolution images of 40–48h APF axons overexpressing *UAS-GFP-Crb*^*WT*^ or *UAS-GFP-Crb*^*RR*^ colocalized with *mtdTomato or mCherry-2xFYVE*. (**L**) Schematic of the role of crb in promoting exuberant branch pruning and the function of crb endocytosis in safeguarding axon integrity. Fisher’s exact test for statistical significance. Scale bar, 5 µm (B, K); 1µm (B, zoom); 20 µm (C). See Figure S5 for additional data.

According to our developmental single-cell RNAseq (scRNA-seq) data^3,5^, c*rb* is expressed in ORNs but not their postsynaptic partner PNs (Figure S5A), indicating that crb in our endocytome is likely produced autonomously by ORNs. Using an endogenously tagged *crb* allele^53^ (*crb-GFP*), we found that crb-GFP proteins appeared punctate and partially colocalized with early endosomes in developing ORN axons (Figure 5B’, B”) with non-colocalized puncta likely falling within other endosomal compartments. Consistent with its moderate endosome enrichment (Figure 4B), some crb signal was not punctate which may be indicative of surface localization (Figure 5B).

Having validated our proteomic identification of crb as an endosome-enriched protein, we next investigated its function in axon development. Pan-ORN knockdown of *crb* caused some DM6-ORN axons to overshoot their target glomerulus and project ectopically into nearby glomeruli (Figure 4G). Using mosaic analysis with a repressible cell marker (MARCM)^54^, we probed the cell-autonomous role for crb using a null allele^55^. We used either *eyeless-FLP* (*ey-FLP*) or *heat shock-FLP* (*hs-FLP*) to induce large (in 30–50% of ORNs) or small (down to single ORN) MARCM clones, respectively, wherein only homozygous *crb* mutant axons expressed a membrane marker in DM6and DL4-ORN axons. Mutant clones displayed significantly increased exuberant branches extending beyond the DM6 glomerulus compared to controls (Figure 5C,D, G and S5B, C, F). This phenotype was rescued by expressing wild-type crb (*UAS-GFP-crb*^*WT*^) only in labeled ORNs (Figure 5E,G and S5D, F), validating that crb acts cell autonomously. During development, DM6 axons send branches to neighboring glomeruli (which are detectable at ∼48h APF) that are later pruned to form the adult circuit^22^ (Figure S5H–J). Their persistence in *crb* mutants suggests that crb promotes the pruning of exuberant branches.

Does endocytosis play a role in crb-dependent pruning? Crb’s highly conserved 37-amino acid cytoplasmic tail contains two lysine (K) residues (Figure 5A) whose ubiquitination drives its endocytosis^56^. We compared the distribution of crb proteins produced from the wild-type crb rescue transgene and an endocytosis deficient version (*UAS-GFP-crb*^*RR*^), which is identical in sequence and genomic insertion site except for two cytoplasmic lysines (K) are mutated to arginines (R). Crb^WT^ was punctate and partially colocalized with an early endosome marker (Figure 5I,J). Crb^RR^ was uniformly distributed across the membrane (Figure 5K) and was largely absent from the early endosomal compartment, consistent impaired internalization

Expression of *crb*^*RR*^ in *crb* mutant ORNs did not alter the number of exuberant branches present in adult DM6 axons (Figure 5F,G and S5E, F), indicating that crb endocytosis is not necessary for axon pruning. Furthermore, developing crb^RR^ axons extended a similar proportion of branches to nearby glomeruli as controls (Figure S5K, L), demonstrating that restricting crb to the plasma membrane does not impair normal features of axon development. Collectively, these results demonstrate that crb at the plasma membrane, not the endosome, is necessary for exuberant branch removal.

Since our data implies that crb promotes developmental axon pruning, we asked if sustained surface crb localization causes over-pruning of mature axons. Strikingly, inhibiting crb endocytosis caused axons to become fragmented (Figure 5F[bottom], H and S5E, G)—hallmarks of axon instability. There were no changes in the number of ORN cell bodies in *crb*^*RR*^ compared to controls (Figure S5M), suggesting that this phenotype is not a consequence of cell death. Thus, prolonged surface localization of crb leads to over-pruning of the entire axon terminal. We propose that the precise balance of crb between endosomes and cell surface enables axon pruning during development while preserving terminal integrity in the mature circuit (Figure 5L). Altogether, these findings highlight how spatially resolved CSP profiling can uncover critical mechanisms that coordinate developmental axon pruning and long-term circuit stability.

### Multi-omics reveals ligand–receptor pairs internalized during axon targeting

Beyond transmembrane proteins, the endocytome contained a diversity of secreted proteins. Because these proteins can originate from neighboring neurons or glia and enter endosomes via receptor-mediated internalization, we asked whether this dataset could be further leveraged to infer cell-cell interactions (Figure 6A).

**Figure 6.**
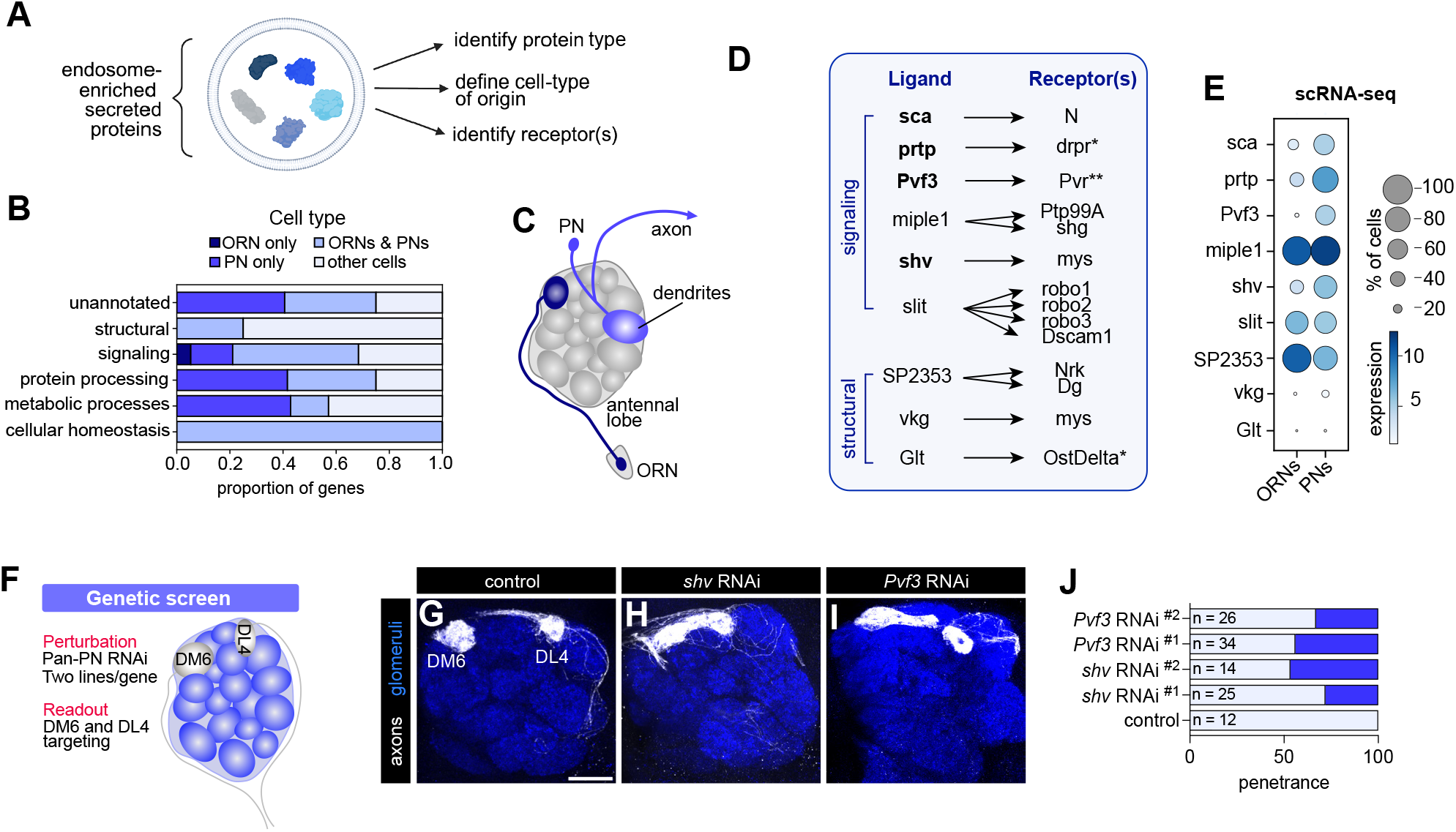
Multi-omic analysis reveals ligand–receptor pairs internalized during axon targeting (**A**) Schematic depicting workflow for identifying endocytosed ligand–receptor pairs. (**B**) Stacked bar plot depicting the cell-type(s) where secreted proteins are robustly expressed (log2[CPM+1] ≥ 4 in ≥ 30% of cells). All endocytome-enriched proteins, including unannotated ones, were used for this analysis. Transcripts not expressed in ORNs or PNs are likely from glia or local interneurons whose transcriptomes have not been profiled. (**C**) Schematic of fly antennal lobe which contains the axons of ∼50 distinct subtypes of ORNs and the dendrites of their ∼50 distinct subtypes of PNs forming one-to-one connection at ∼50 glomeruli. (**D**) Potential ligand–receptor pairs found in ORN endosomes. These pairs were identified based on data found on Flybase (www.flybase.org). Bolded ligands are more abundantly expressed in PNs. *, endosome-enriched; **, similarly labeled in endocytome and surfaceome. (**E**) Dotplot depicting expression levels of transcripts encoding ligands in (D) in ORNs or PNs. scRNA-seq expression is log_2_(CPM+1). CPM = counts per million reads. Dotplot depicts averaged expression in all PN or ORN types at 24h and 48h APF. (**F**) Schematic depicting loss-of-function genetic screen where secreted proteins were knocked-down in all PNs (via *VT033006-GAL4)* and axons projecting to the DM6 and DL4 glomeruli were labeled with *AM29-QF2* driven membrane-targeted tdTomato (*QUAS-mtdTomato*; white). Scale bar is 20 µm. (**G**–**I**) Representative images of antennal lobes depicting phenotypes resulting from knockdown of each secreted protein. (**J**) Quantification of phenotypic penetrance for DM6-axons in indicated genotypes. See Figure S6, 7 for additional data.

The 86 endosome-enriched secreted proteins (Table S2) spanned five functional categories: (1) cellular homeostasis; (2) protein processing; (3) signaling; (4) metabolic processes; and (5) structural roles (Figure 6B,S6A). The cellular homeostasis category included the complete ferritin storage complex, and collagens (e.g., vkg, Col4a1) and laminin (e.g., wb) were within the structural group (Figure S6A), suggesting roles for iron and extracellular matrix uptake in neuronal development. Proteins within the metabolic process and protein processing categories were responsible for regulating extracellular breakdown of macromolecules as well as growth factor and hormone signaling, respectively (Figure S6A). Signaling proteins were the largest category, containing both classic neurodevelopment regulators (e.g., slit [sli], and scabrous [sca]) and proteins not previously linked to development (Figure S6A). These findings highlight the rich extracellular milieu axons are navigating.

To begin decoding cell-cell communication underlying axon development, we next asked which cell types secreted the proteins internalized by ORNs. Using our developmental scRNA-seq data^3,5^, we mapped the expression of each protein to ORNs or PNs (Figure 6C; see STAR Methods for details). Many transcripts were shared between ORNs and PNs (Figure 6B, S7). Although only two transcripts were abundantly expressed in ORNs, over 35% of transcripts in the unannotated, protein processing, and metabolic process categories, as well as 15% of signaling transcripts, were more abundant in PNs (Figure 6B). To corroborate these cell-type-assignments, we experimentally validated PN-specific expression of two such transcripts, *Pxn* and *sca* (Figure S6B–D). Together, these data indicate that endocytome profiling is sensitive enough to detect cues secreted by neighboring cells and endocytosed by ORN axons.

We next evaluated whether any of the known receptors for these secreted proteins were present in endosomes. Indeed, receptors for several signaling and structural proteins were in endosomes (Figure 6D). Although not all receptors were endosome-enriched (Figure 6D), these data nevertheless support ligand entry via receptor binding and internalization. Lastly, we tested whether any of the ligands are secreted by PNs to regulate ORN axon targeting (Figure 6F,J). We focused on *Pvf3* and *shv*, as both were more abundant in PNs (Figure 6E) and neither has been linked to olfactory circuit development. Knockdown of *shv* (encoding a soluble ß-integrin [mys] ligand) in all PNs caused DM6-ORNs to mistarget laterally and at times fuse with DL4 axons (Figure 6H,J). Whereas knockdown of *Pvf3* (see below) led to random mistargeting of DM6 axons and widespread glomerular disorganization, including fusion of some glomeruli and loss of others, consistent with a global disruption of circuit assembly (Figure 6I,J). Altogether, these analyses highlight how cell-type-specific endosome profiling reveals intercellular signals that shape tissue development and organization.

### Endocytosis tunes Pvr levels to match target-derived Pvf3 for axon targeting

Finally, we investigated how the internalization of one of the ligand–receptor pairs we discovered, Pvf3–Pvr, contributes to axon target selection. Pvf3 is one of three fly orthologs of both platelet-derived (PDGF) and vascular endothelial growth factor (VEGF)^57,58^, and the only one expressed in PNs (Figure S8A). Pvr is a receptor tyrosine kinase (Figure 7A) and the sole fly ortholog of the PDGF and VEGF receptors^57^. This pair has been shown to regulate phagocyte migration^59^, but their role in neural development is undefined.

**Figure 7.**
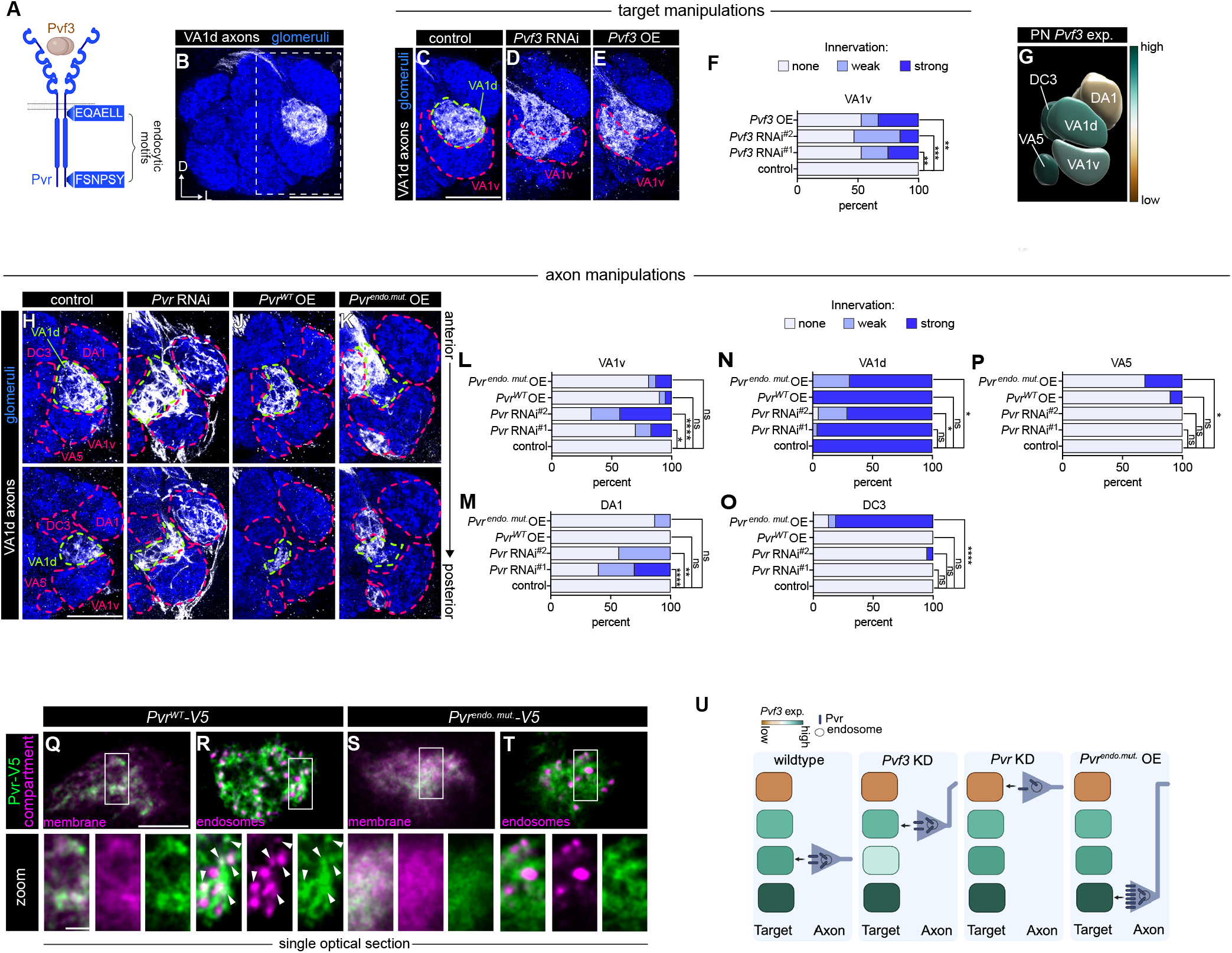
Endocytosis tunes Pvr levels to match target-derived Pvf3 for axon targeting. (**A**) Schematic of Pvf3 and Pvr. Boxes indicate endocytic motifs and their approximate location flanking the tyrosine kinase domain. (**B**) Image of the antennal lobe depicting VA1d-ORN axons. Dashed rectangle indicates glomeruli that are depicted in subsequent panels. (**C**–**E**) Images of antennal lobes depicting targeting of VA1d-ORN axons when *Pvf3* expression is manipulated in PNs (left) and morphology of glomeruli analyzed (right). (**F**) Quantification of VA1d-ORN axon targeting when *Pvf3* is manipulated in postsynaptic targets. n = 21 (controls); n = 26 (*Pvf3 RNAi*^*#1*^); n = 36 (*Pvf3 RNAi*^*#2*^); n = 21 (*Pvf3* overexpression [OE]). (**G**) Volume rendering depicting *Pvf3* expression in glomeruli analyzed in prior and subsequent panels. Data used to generate expression heatmap overlay is in Figure S6B–E. Expression is in arbitrary units. (**H**–**K**) Images of the antennal lobe depicting targeting of VA1d-ORN axons when *Pvr* is manipulated. (**L**–**P**) Quantification of VA1d axon targeting when *Pvr* is manipulated in ORN axons. n = 22 (control); n = 23 (*Pvr RNAi*^*#1*^); n = 21 (*Pvr RNAi*^*#2*^); n = 23 (*Pvr*^*WT*^ OE); n = 16 (*Pvr*^*endo*.*mut*.^ OE). (**Q**–**T**) Airyscan super-resolution images of 40–48h APF axons overexpressing *UAS-Pvr*^*WT*^*-V5* or *UAS-Pvr*^*endo*.*mut*.^*-V5* along with *mtdTomato or mCherry-2xFYVE*. Arrowheads denote Pvr puncta colocalized with endosomes. (**U**) Schematic of the role of Pvf3-Pvr in axon targeting. Fisher’s exact test was used to determine statistical significance. Scale bar: 20 µm (B, C, H); 5 µm (Q, top), 2 µm (Q, bottom). See Figure S8–10 for additional data.

We first sought to comprehensively characterize *Pvf3*’s expression in PNs to better understand how its loss caused such wide-spread disruptions in antennal lobe morphology (Figure 6I). Analysis of scRNA-seq revealed differential expression of *Pvf3* between PN types (Figure S8A); however, not all PN types are present in this dataset. To validate and extend this analysis, we used an intersectional *GAL4-*based reporter strategy which largely mirrored the *Pvf3* expression pattern observed in our PN transcriptomes (Figure S8B, C). The marked heterogeneity in *Pvf3* expression across PN types led us to hypothesize that cell-type-specific ligand levels instruct ORN axon targeting.

Having genetic access to both VA1d-PNs and VA1d-ORNs during development enabled us to investigate the role of PN-derived Pvf3 in regulating ORN axon targeting to the VA1d glomerulus (Figure 7B). *Pvf3* is expressed at a relatively high level in VA1d-PNs (Figure 7G and S8C). *Pvf3* knockdown in VA1d-PNs (and DC3/DA1-PNs) caused VA1d-ORNs to mistarget to the neighboring VA1v glomerulus (Figure 7C, D, F). Indicating that Pvf3 is secreted by PNs to regulate ORN axon targeting, pan-ORN *Pvf3* RNAi did not cause significant targeting defects (Figure S9A). Intriguingly, overexpression of *Pvf3* in VA1d-PNs (and DC3/DA1-PNs) also caused VA1d-ORN axons to mistarget to the VA1v glomerulus (Figure 7E,F). Among VA1d’s neighbors, VA1v-PNs exhibited the most similar *Pvf3* expression to VA1d-PNs, albeit at a slightly lower level (Figure 7G). One hypothesis that could account for these observations is that Pvr directs ORN axons to glomeruli that express Pvf3 at a level most comparable to their original targets.

If Pvr mediates ORN responsiveness to Pvf3, then altering receptor levels should affect axon targeting. Indeed, VA1d-ORN-specific knockdown of *Pvr* caused increased targeting to the DA1 and VA1v glomeruli (Figure 7H, I, L, M). Both glomeruli express lower levels of *Pvf3* than VA1d (Figure 7G), implying that when receptor levels are reduced, ORN axons target glomeruli with lower ligand expression. We therefore hypothesized that *Pvr* overexpression would enable VA1d-ORNs to target glomeruli where ligand levels are higher than their original target. Thus, we generated a *Pvr* overexpression transgene (*UAS-Pvr*^*WT*^). However, elevated *Pvr* levels in VA1d-ORNs did not significantly alter axon target selection (Figure 7J, L–P).

We hypothesized that ORNs may actively downregulate excess Pvr via endocytosis to maintain the receptor levels needed to match with target-derived Pvf3.

To test this, we identified two putative endocytic motifs flanking Pvr’s tyrosine kinase domain (Figure 7A) and generated a mutant transgene, identical in sequence and genomic location to *UAS-Pvr*^*WT*^, with these residues mutated to alanines (*UAS-Pvr*^*endo*.*mut*.^). The tyrosine (Y) residue in the second motif was left intact to minimally disrupt signaling. When expressed in ORNs, both Pvr^WT^ and Pvr^endo.mut.^ trafficked to axon terminals (Figure 7Q,S). Consistent with the endocytome data where Pvr is equally distributed between the surface and endosomes (Figure 6D), Pvr^WT^ was present in both compartments (Figure 7Q,R). By contrast, Pvr^endo.mut.^ appeared mostly at the cell surface and did not strongly colocalize with endosomes (Figure 7S,T), a distribution also observed in cultured cells (Figure S10).

Interestingly, overexpression of endocytosis-deficient *Pvr* caused VA1d-ORN axons to mistarget to DC3 and VA5 glomeruli (Figure 7K,N–P). Consistent with the matching-level hypothesis, both DC3 and VA5 glomeruli express *Pvf3* at higher levels than VA1d (Figure 7G). Finally, we probed the broader requirement for this signaling mechanism in DM6-ORNs and could recapitulate all *Pvr* loss- and gain-of-function phenotypes in these neurons (Figure S9B–G). Together, these data indicate that endocytosis finely tunes axonal Pvr surface presentation to match Pvf3 levels in the target (Figure 7U). Altogether, these data reveal how developing neurons use endocytosis to actively modulate CSP localization and control axon target selection.

## DISCUSSION

Here, we developed endocytome profiling as a systematic approach to decode how the cell-surface proteome is dynamically remodeled across cell states, developmental stages, and disease contexts. We used this method to expose the scope of membrane remodeling in developing axons and to unveil how it enables neurons to adjust their surface proteome to meet stage-specific needs. Below, we highlight the biological insights unlocked by this technological advance and discuss the avenues it opens for future discovery.

### Endocytome profiling unlocks new dimensions of proteome remodeling in intact tissues

Proximity labeling-based has been used to define: 1) how the surface proteome differs between distinct neural cell types^60^, 2) the neuron-astrocyte proteomic interface^61^, and 3) steady-state proteome landscape changes over developmental time^6,7^. Endocytome profiling achieves these goals and offers three key innovations.

First, it transforms static snapshots of the surface proteome into a dynamic view of its remodeling at any given time *in situ*. Notably, our proteomes are derived from axons from a specific cell type constituting less than 5% of the total axons in the fly brain—a major advance as such remodeling has, until now, been studied primarily in culture systems^8–10^. Second, beyond capturing CSP dynamics, this approach also reveals trans-cellular communication by detecting proteins internalized from neighboring cells. Finally, it enables high-quality profiling of the endolysosomal network from sparse populations, overcoming potential limitations of other methods which have thus far been restricted to bulk neural populations^39,62,63^. This paves the way for uncovering how membrane trafficking varies across cell states and more broadly demonstrates that proximity labeling can be used for organelle-specific proteomics with cell-type resolution.

As was the case with cell surface proteomics approaches in mammals^7,60^, which we first established in *Drosophila*^6^, we expect that endocytome profiling will be readily adapted for use in other model systems. Broad application of this tool is poised to reveal how the surface proteome is reconfigured to support activity-dependent morphological plasticity, to identify proteins transcytosed across the blood-brain barrier, to uncover trafficking defects in disease models with endolysosomal dysfunction, and to explore a range of biological processes both within and beyond the nervous system. This positions endocytome profiling as a powerful approach for dissecting cell-surface remodeling and membrane trafficking across diverse biological and pathological contexts.

### The endocytome reveals CSP regulation underlying neuronal connectivity

Endocytome profiling provided a systems-level view of how endocytosis actively sculpts the neuronal surface proteome to regulate circuit connectivity. Although this has been appreciated at the single-CSP level^14–16,64^, our mechanistic studies elucidated that even within a single ORN type (DM6-ORNs) and timepoint, endocytosis regulates multiple CSPs to coordinate the developmental events necessary to build and maintain neural circuits.

This work revealed that developing axons may repurpose cell-cell junction proteins as part of a conserved developmental program. Accordingly, a recent study found that transcripts encoding junctional proteins are enriched in central mammalian neurons during neurite outgrowth^4^ and a *C. elegans* tight junction CSP regulates dendrite growth^65^. In line with this, nearly all endosome-enriched junctional proteins had either direct mammalian orthologs or were part of conserved families present across phyla (Table S3). Our loss-of-function screen demonstrated that this class of CSPs broadly regulates axon development. Altogether, these findings indicate that junctional proteins exhibit conserved neuronal functions and implies that their roles in circuit development are actively shaped by endocytic regulation.

As an example of such regulation, our analysis of crb uncovered both a new neuronal function for this junctional protein and showed how endocytosis sculpts the surface proteome to meet stage-specific developmental needs. Specifically, crb promotes axon pruning during development but must be downregulated via endocytosis to prevent over-pruning of the mature terminal (Figure 5L). Sustained surface localization of crb destabilizes ORN axons, which contrasts its role in photoreceptors where *crb* loss leads to degeneration^49^ and *CRB1* and *CRB2* mutations are linked to retinal dystrophies in humans^50^. Thus, while crb is essential for photoreceptor integrity it must be tightly regulated in ORNs to preserve axon stability. Given its conserved retinal functions and the expression of *CRB1*/*CRB2* in the developing mammalian brain^51,52^, it is possible that these orthologs also mediate neurite pruning in mammals. Finally, the genetic tractability of *Drosophila* provides an excellent platform to dissect the mechanisms of crb-mediated pruning, which may reveal additional conserved functions of this protein.

A strength of cell type-specific profiling is the ability to interrogate cell-cell interactions within intact tissues. Our analysis demonstrated that endocytome data can be integrated with complementary transcriptome profiling approaches to reveal transcellular signaling events. In so doing, we discovered that an endosome-enriched growth factor Pvf3 is secreted by postsynaptic partner PNs to regulate ORN axon target selection via its receptor Pvr. Though *Pvf3* is expressed in most PN types, its differential expression is required for ORN axon targeting, raising the question of how a broadly expressed ligand can mediate ORN-type specific outcomes. PDGF, Pvf3’s vertebrate ortholog, triggers a concentration-dependent switch between migration and proliferation by activating distinct signaling pathways downstream of the PDGF receptor^66^. ORN axons transiently explore neighboring glomeruli before stabilizing in their target^22,67^. Thus, varying postsynaptic Pvf3 levels could trigger different signaling responses in distinct ORN types, promoting retraction from incorrect glomeruli and stabilization in the correct target. Future work addressing the signaling outcomes downstream of Pvf3-Pvr activation will be important for fully understanding the role of these proteins in circuit assembly.

In summary, endocytome profiling provides a versatile framework for mapping surface proteome dynamics across cell types, developmental stages, and species. Our findings demonstrate how profiling CSP distribution in the native tissue environment reveals key principles of neuronal development and cellular organization. Broad application of this approach should not only expand our understanding of how the plasma membrane protein landscape is remodeled in health and disease but also uncover generalizable mechanisms of cellular plasticity.

### Limitations of this study

The limitations of HRP-based proximity labeling have been described elsewhere^68^; here we highlight considerations specific to our approach. First, while HRP^endo^ has high specificity for the endolysosomal compartment, it is not restricted to a distinct subset of vesicles, such as recycling endosomes or lysosomes. Thus, follow-up studies are needed to determine the trafficking fate(s) of internalized cargos. Second, a minimum number of cells per brain is likely required to express HRP^endo^ to generate robust signal for mass-spectrometry. Consequently, our proteomes are from all ORNs rather than distinct subtypes, which may obscure low-abundance or subtype-specific cargos. It is possible that utilizing labeling substrates not naturally found in samples such as alkyne-phenol^69^ (in lieu of biotin-phenol) could improve the signal-to-noise ratio and enable profiling of lower-abundance cell types. Nevertheless, we anticipate that the increasing sensitivity of mass-spectrometry will enable subtype-specific profiling in the future.

## ACKNOWLEDGEMENTS

We thank members of the Luo lab, especially D. Pederick, A. Starr, T. Hindmarsh Sten, and Y. Ge for support, insight, and feedback on this study. We also thank K. Eichel, K. Shen, C. Taylor, J. Vaughn for feedback on this study. We are grateful to M. Pellikka and U. Tepass for insight into crumbs biology and reagents. R. Ward, Y. Hong, Addgene, Bloomington *Drosophila* Stock Center, Vienna *Drosophila* Stock Center, and Best Gene, provided critical reagents. We appreciate the administrative assistance from M. Molacavage. C.N.M. was a HHMI fellow of the Damon Runyon Cancer Research Foundation (DR-2390-20). J.L is a group leader of the HHMI Janelia Research Campus. L.L is a HHMI investigator. This work was supported by the National Institutes of Health (R01-DC005982 to L.L. and K99-DC021195 to C.N.M.) and Wu Tsai Neurosciences Institute of Stanford University (A.Y.T. and L.L.).

## AUTHOR CONTRIBUTIONS

C.N.M. and L.L. conceived this project. C.N.M. a L.L. designed the proteomic experiments with inp from A.Y.T., W.Q., S.A.C., N.D.U., and C.X. C.N.M., C.X. C.L., Z.L., K.L.W., D.J.L, K.X.D, H.J. dissected fly brains for the proteomic experiment. C.N.M. processed samples for proteomics with advice from C.X., N.D.U., C.X., S.A.C. performed post-enrichment sample processing, mass spectrometry, and initial data analysis. C.N.M. analyzed proteomic data with input from J. L. H.J. and K.X.D. performed co-localization experiments and dissections for phenotypic analysis. D.J.L. produced transgenic flies. C.N.M. performed and analyzed all other experiments with input from L.L and assistance from K.X.D. and H.J. C.N.M. wrote the manuscript with input from L.L. and all-coauthors. L.L. supervised the work.

## RESOURCE AVAILABILITY

### Materials availability

We will deposit newly generated constructs in Addgene, and newly generated transgenic flies in the Bloomington Drosophila Stock Center. All other unique reagents generated in this study are available from the lead contact.

### Data and code availability

The original mass spectra and the protein sequence databases used for searches have been deposited in the public proteomics repository MassIVE (http://massive.ucsd.edu) and are accessible at ftp://MSV000097831@massive-ftp.ucsd.edu when providing the dataset password: endocytome. If requested, also provide the username: MSV000097831. These datasets will be made public upon acceptance of the manuscript. Processed proteomic data is provided in Table S2.

## EXPERIMENTAL MODEL

### Drosophila stocks and genotypes

Flies were maintained on standard cornmeal media with a 12 hr light-dark cycle at 25ºC, except for RNAi/overexpression crosses which were raised at 29ºC. Complete genotypes for flies used in each experiment are described in Table S1.

We note that *Peb-GAL4*, the driver used to express UAS-HRP transgenes, is present in neurons in the visual and auditory systems as well as ∼20 other neurons in the brain. Our dissections removed the visual system but not the other neurons. However, they comprise less than 10% of the axons profiled and thus represent a minor fraction of our proteomes.

## METHOD DETAILS

### Generation of UAS constructs and transgenic flies

*UAS-HA-HRP*^*surf*^ and *UAS-HA-HRP*^*endo*^ flies were generated from a gBlock containing a wingless signal peptide upstream of an epitope tag (HA) followed by HRP fused to the N terminal regions of human CD2 The gBlock was cloned into a 5x *pUASt-attB* vector. The cytoplasmic tail of CD2 was mutated via site-directed mutagenesis (New England Biolabs) to include a dileucine endocytic motif or a motif consisting of all alanines. Aside from the addition of the HA tag and dileucine motif (or alanines), these transgenes are otherwise identical to the one used in Li et al., 2020. The constructs were validated by full-length plasmid sequencing and injected into embryos bearing the *attP24* landing site. G0 flies were crossed to a white– balancer, and all white+ progeny were individually balanced.

*UAS-Pvf3-myc, UAS-Pvr*^*WT*^*-V5, UAS-Pvr*^*endo*.*mut*.^*-V5, UAS-2xFYVE-mCherry* flies were synthesized and cloned by Twist Biosciences into a 10x *pUASt-attB* vector. cDNA sequences for the isoforms of Pvf3 (isoform D) and Pvr (isoform J) identified in our proteomics were used to generate each rescue transgene. Epitope tags are C-terminal just prior to the stop codon. 2xFYVE-mCherry was fly codon optimized. The constructs were all validated full-length plasmid sequencing and injected into embryos bearing an attP40 (*UAS-2xFYVE-mCherry*,), attP2 (*UAS-Pvf3-myc*), or attP86Fb (*UAS-Pvr* constructs, *UAS-2xFYVE-mCherry*). G0 flies were crossed to a white– balancer and all white+ progeny were individually balanced.

All flies were injected in-house using standard microinjection methods, except *UAS-Pvr*^*WT*^*-V5* which was made through Best Gene Inc.

### HRP-mediated proximity biotinylation of proteins in ORN endosomes and surface

Proximity labeling was performed using previously published methods^6^ with one key difference in the usage of the membrane-permeable biotin-phenol instead of membrane-impermeable BXXP as a substrate which enables the labeling of proteins in the intracellular compartments in addition to just extracellular or surface proteins. Briefly, 30–36h APF (after puparium formation) brains containing ORN undergoing axon target selection were dissected (optic lobes were removed) in ice cold Schneider’s media (ThermoFisher) and transferred into a 1.5 mL protein low-binding tube (Eppendorf) containing additional media on ice. Brains were washed with Schneider’s media to remove debris and incubated in 500 µM biotin-phenol (BP; APExBio) in Schneider’s media while rotating at 4ºC for 1 hr. Brains were subsequently labeled with 1 mM (0.003%) H_2_O_2_ (ThermoFisher) for 10 min while rotating and immediately quenched by five thorough washes using the quenching buffer (10 mM sodium ascorbate, 5 mM Trolox, and 10 mM sodium azide in phosphate buffered saline [PBS]). Following these washes, the quenching solution was removed, and brains were either fixed for immunostaining (see below for details) or were snap frozen and stored at –80ºC for downstream processing for proteomic analysis.

### Enrichment of biotinylated proteins

Brains were processed in the original collection tube to avoid loss during transferring using previously published methods^6,67,87^. Briefly, 40 µL of high-SDS RIPA buffer (50 mM Tris-HCl [pH 8.0], 150 mM NaCl, 1% sodium dodecyl sulfate [SDS], 0.5% sodium deoxycholate, 1% Triton X-100, 1x protease inhibitor cocktail [Sigma-Aldrich], and 1 mM phenylmethylsulfonyl fluoride [PMSF; Sigma-Aldrich]) was added to each tube and frozen brains were homogenized on ice. Next, samples of the same experimental group were spun down and merged, rinsed with additional 100 µL of high-SDS RIPA, vortexed, and sonicated briefly. Lysates were diluted with 1.2 mL of SDS-free RIPA buffer (50 mM Tris-HCl [pH 8.0], 150 mM NaCl, 0.5% sodium deoxycholate, 1% Triton X-100, 1x protease inhibitor cocktail, and 1 mM PMSF) and rotated for 1h at 4ºC. Lysates were then diluted with 200 µL of normal RIPA buffer (50 mM Tris-HCl [pH 8.0], 150 mM NaCl, 0.2% SDS, 0.5% sodium deoxycholate, 1% Triton X-100, 1x protease inhibitor cocktail, and 1 mM PMSF), transferred to a 3.5 mL ultracentrifuge tube (Beckmann Coulter), and centrifuged at 100,000 × g for 30 min at 4ºC. 1.5 mL of supernatant was collected for each sample and added to 210 µL of pre-washed streptavidin magnetic beads (Pierce) and incubated while rotating at 4ºC overnight. The next day, beads were washed twice with 1mL RIPA buffer, once with 1 mL of KCl, then with 1mL 0.1 M Na_2_CO_3_, followed by 1 mL 2 M urea (in 10 mM Tris-HCl [pH 8.0]), and finally twice with 1 mL RIPA buffer.

Finally, beads were washed twice in 1 mL NaCl (75 mM NaCl in 50 mM Tris HCl [pH 8]) buffer and resuspended in 400 µL of NaCl buffer. 10% of the bead suspension was removed for Western blot analysis, and the rest were snap frozen prior to on-bead digestion.

#### Western blotting of biotinylated proteins

Biotinylated proteins were eluted from streptavidin beads via the addition of 20 µL of elution buffer (2X Laemmli sample buffer [BioRad], 20 mM dithiothreitol [Sigma-Aldrich], and 2mM biotin [Sigma-Aldrich]) followed by a 10 min incubation at 95ºC. Proteins were loaded on to 4%–12% Bis-Tris PAGE gels (ThermoFisher) and transferred to PVDF membranes (ThermoFisher). After blocking with Intercept (TBS) blocking buffer (LI-COR) for 1h, membranes were incubated with Streptavidin-800 (LI-COR) and visualized using the ChemiDoc imaging system (BioRad).

#### On-bead trypsin digestion of biotinylated proteins

Peptides bound to streptavidin magnetic beads were washed four times with 200 µL of 50 mM Tris-HCl (pH=7.5) buffer. After removing the final wash, the beads were incubated twice at room temperature (RT) in 80 µL of the digestion buffer – 2 M Urea, 50 mM Tris-HCl, 1 mM DTT, and 0.4 µg trypsin – while shaking at 1000 rpm. The first incubation lasted 1 hour, followed by the second incubation of 30 minutes. After each incubation, the supernatant was collected and transferred into a separate tube. The beads were then washed twice with 60 µL of 2 M Urea/50 mM Tris-HCl buffer. The resulting washes were combined with the digestion supernatant. The pooled eluate of each sample was then spun down at 5000 × g for 30 sec to collect the supernatant. The samples were subsequently reduced with 4 mM DTT for 30 minutes at RT with shaking at 1000 rpm, followed by alkylation with 10 mM Iodoacetamide for 45 min in the dark at RT while shaking at 1000 rpm. Overnight digestion of the samples was performed by adding 0.5 µg of trypsin to each sample. The following morning, the samples were acidified with neat formic acid (FA) to the final concentration of 1% FA (pH<3).

Digested peptide samples were desalted using in-house packed C18 (3M) StageTips. C18 StageTips were conditioned sequentially with 100 µL of 100% methanol (MeOH), 100 µL of 50% (vol/vol) acetonitrile (MeCN) with 0.1% (vol/vol) FA, and two washes of 100 µL of 0.1% (vol/vol) FA. Acidified peptides were loaded onto the C18 StageTips and washed twice with 100 µL of 0.1% FA. The peptides were then eluted from the C18 resin using 50 µL of 50% MeCN/0.1% FA. The desalted peptide samples were snap-frozen and vacuum-centrifuged until completely dry.

### TMT labeling and stagetip peptide fractionation

Desalted peptides were labeled with TMT16 reagents (Thermo Fisher Scientific). For this experiment, relevant TMT channels are 131N-ORN_NC 1, 132C-ORN_NC 2, 128C-ORN_HRP^endo^ 1, 133N-ORN_ HRP^endo^ 2, 133C-ORN_ HRP^endo^ 3, 129C-ORN_ HRP^surf^ 1, 130N-ORN_ HRP^surf^ 2, 127C-ORN_ HRP^surf^ 3. Each peptide sample was resuspended in 80 μL of 50 mM HEPES and labeled with 20 µL of the 25 µg/µL TMT reagents in MeCN. The samples were then incubated at RT for 1 hour while shaking at 1000 rpm. To quench the TMT-labeling reaction, 4 μL of 5% hydroxylamine was added to each sample, followed by a 15-minute incubation at RT with shaking. TMT-labeled samples were combined and vacuum-centrifuged to dry. The samples were then reconstituted in 200 μL of 0.1% FA and desalted on a C18 StageTip using the previously described protocol. The desalted TMT-labeled combined sample was then dried to completion.

The combined TMT-labeled peptide sample was fractionated by basic reverse phase (bRP) fractionation using an in-house packed SDB-RPS (3M) StageTip. A StageTip containing three plugs of SDB material was prepared and conditioned with 100 μL of 100% MeOH, 100 μL of 50% MeCN/0.1% FA, and 2x with 100 μL of 0.1% FA. The sample was resuspended in 200 µL 0.1% FA (pH < 3) and loaded onto the conditioned StageTip and eluted in a series of buffers with increasing MeCN concentrations. Eight fractions were collected in 20 mM ammonium formate (5%, 7.5%, 10%,,12.5%, 15%, 20%, 25%, and 45% MeCN), dried to completion and analyzed by LC-MS/MS.

### Liquid chromatography and mass spectrometry

All peptide samples were separated and analyzed on an online liquid chromatography tandem mass spectrometry (LC-MS/MS) system, consisting of a Vanquish Neo UPHLC (Thermo Fisher Scientific) coupled to an Orbitrap Exploris 480 (Thermo Fisher Scientific). All peptide fractions were reconstituted in 9 µL of 3% MeCN/0.1% FA. 4 µL of each fraction was injected onto a microcapillary column (Picofrit with 10 µm tip opening/75 µm diameter, New Objective, PF360-75-10-N-5), packed in-house with 30 cm of C18 silica material (1.5 µm ReproSil-Pur C18-AQ medium, Dr. Maisch GmbH, r119.aq) and heated to 50 °C using column heater sleeves (PhoenixST). Peptides were eluted into the Orbitrap Exploris 480 at a flow rate of 200 nL/min. The bRP fractions were run on a 154 min-method. Solvent A comprised 3% acetonitrile/0.1% FA. Solvent B comprised 90% acetonitrile/0.1% FA. The LC-MS/MS method used the following gradient profile: (min: %B) 0:2; 1:6; 122:35; 130:60; 133:90; 143:90; 144:50; 154:50 (the last two steps at 500 nl/min flow rate).

Mass spectrometry was conducted using a data-dependent acquisition mode, MS1 spectra were measured with a resolution of 60,000, a normalized AGC target of 100%, and a mass range from 350 to 1800 m/z. MS2 spectra were acquired for the top 20 most abundant ions per cycle at a resolution of 45,000, an AGC target of 50%, an isolation window of 0.7 m/z and a normalized collision energy of 32. The dynamic exclusion time was set to 20 s.

### Mass spectrometry data processing

Mass spectrometry data was processed using Spectrum Mill (proteomics.broadinstitute.org). Spectra within a precursor mass range of 600-6000 Da with a minimum MS1 signal-to-noise ratio of 25 were retained. Additionally, MS1 spectra within a retention time range of ± 45 s, or within a precursor m/z tolerance of ± 1.4 m/z were merged. MS/MS searching was performed against a human Uniprot database. For searching, fixed modifications were TMT16-Full-Lys modification and carbamidomethylation on cysteine. Variable modifications included acetylation of the protein N-terminus, oxidation of methionine and cyclization to pyroglutamic acid. Digestion parameters were set to “trypsin allow P” with an allowance of 4 missed cleavages. The matching tolerances were set with a minimum matched peak intensity of 30%, precursor and product mass tolerance of ± 20 ppm.

### Quantitative comparison of endosome and surface proteomes

Peptide spectrum matches (PSMs) were validated with a maximum false discovery rate (FDR) threshold of 1.2% for precursor charges ranging from +2 to +6. A target protein score of 9 was applied during protein polishing auto-validation to further filter PSMs. TMT16 reporter ion intensities were corrected for isotopic impurities using the afRICA correction method in the Spectrum Mill protein/peptide summary module, which utilizes determinant calculations according to Cramer’s Rule. Protein quantification and statistical analysis were performed using the Proteomics Toolset for Integrative Data Analysis (Protigy, v1.0.7, Broad Institute, https://github.com/broadinstitute/protigy). Differential protein expression was evaluated using moderated t-tests, with P-values calculated to assess significance.

### Proteomic data cutoff analysis

We used a ratiometric strategy^28^ to remove contaminants. Briefly, all detected proteins were annotated as either true-positives (TPs; proteins with a signal peptide and/or a transmembrane domain) or false-positives (FPs; those without either domain) according to the UniProt database. For each experimental group, we calculated the TMT ratios of proteins within this group compared to an averaged negative control value and sorted proteins in descending order. For each TMT ratio, a true-positive rate (TPR) and false positive rate (FPR) were calculated by adding the number of TPs or FPS with a higher ranking and dividing them by the total number of TPs and FPs, respectively. The TPRs and FPRs were used to generate the ROC curve. Cutoffs for experimental groups were determined by finding the TMT ratio where [TPR – FPR] is maximized. Proteins with TMT ratios higher than the cutoff in each experimental group were retained. Finally, proteins present in two of three experimental groups were retained for downstream analysis.

### Immunostaining

Fly brains were dissected according to a previously published protocol^88^. In brief, brains were dissected in PBS, transferred to a tube containing 4% paraformaldehyde in PBST (0.3% Triton X-100), and fixed for 20 min while nutating at RT. Following fixation, brains were washed 3 times for 20 minutes in PBST and blocked for at least 30 min in PBST + 5% normal donkey serum. The following antibodies were used: rat anti-Ncad (Developmental Studies Hybridoma Bank; 1:40), chicken anti-GFP (Aves Labs; 1:1000), rabbit anti-HA (Cell Signaling Technologies; 1:200), rat anti-HA (Sigma Aldrich; 1:200), mouse anti-V5 (ThermoFisher Scientific; 1:100), guinea pig anti-Mcr^70^; (1:500), rabbit anti-dsRed (Takara Bio; 1:1000), rabbit anti-Lamp1 (Abcam; 1:100), rabbit anti-Rab5 (Abcam; 1:100), mouse anti-Rab7 (Developmental Studies Hybridoma Bank; 1:10), goat anti-HRP-594 (Jackson ImmunoResearch; 1:300), mouse anti-mCherry (ThermoFisher Scientific; 1:1000), and rat anti-V5 (Abcam; 1:500) and incubated with brains in block buffer overnight at 4ºC while nutating. Brains were subsequently washed three times for 20 min in PBST and incubated in secondary antibodies (Alexa Fluor 488; Alexa Fluor 564; Alexa Fluor 647; 1:200) overnight at 4ºC while nutating. Brains were again washed three times for 20 min in PBST, transferred to SlowFade antifade reagent (ThermoFisher) and stored at 4ºC prior to mounting. Neutravidin-647 (ThermoFisher) was used to visualize biotinylated proteins.

### Image acquisition and processing

Images were obtained on a Zeiss LSM900 laser-scanning confocal microscope (Carl Zeiss) using either a 40x oil immersion objective (axon targeting experiments) or a 63x oil immersion objective (Airyscan experiments). 16-bit z-stacks for axon targeting experiments were acquired at 1 µm intervals at a resolution of 1024 × 1024. Airyscan images were taken at software optimized resolution and intervals. Brightness and contrast adjustments as well as image cropping was done using Photoshop or Illustrator (Adobe).

### RNAi-based genetic screen

The AM29 screening line was generated by converting *AM29-GAL4* to *AM29-QF2* via the homology assisted CRISPR knock-in (HACK) method^89^. *AM29-QF2* was recombined with *QUAS-mtdTomato* on the second chromosome and put with *Peb-GAL4, UAS-dcr2* on the X chromosome. Virgin females from the AM29 screening line were crossed to *UAS-RNAi* males and the progeny were kept at 25ºC for 2–5 days following egg laying and then transferred to 29ºC to enhance transgene expression. Brains were dissected, processed, and imaged as described above. *Peb-GAL4* is expressed in some peripheral tissues and expression of some RNAi lines using this driver caused adult lethality. To circumvent this, we dissected late-stage pupae of these genotypes and evaluated their axon targeting.

For this analysis, we identified glomeruli using NCad labeling (based on the stereotypy of their size, shape, and positions), and categorized the extent of innervation into each glomerulus (not innervated; weakly innervated; strongly innervated). Axon targeting analysis was performed blinded to genotype when possible. Data was analyzed using Prism10 (GraphPad). For each glomerulus, we calculated the frequency of each type of innervation and plotted the results as stacked bar charts. Fisher’s exact test was performed on innervation frequencies in each glomerulus to determine statistical significance compared to controls. P-values were adjusted using the Bonferroni correction.

### shibire^ts^ temperature shift paradigm

*AM29-GAL4* virgins were crossed to males bearing *UAS-shi*^*ts*^ and allowed to reproduce at room temperature for 2–5 days. Since rearing flies at temperatures different from 25ºC (standard rearing temperature) affects developmental progression, we interpolated the developmental timepoint of flies at each temperature (18ºC or 30ºC) using published developmental progression data^90^. Progeny were moved to 18ºC (restrictive temperature) until pupariation. Once pupae formed, 0–6 h pupae were collected, and allowed to develop until they were ∼24h APF. At this time, flies were moved to a 30ºC incubator for 24 h. Pupae were then returned to 18ºC and allowed to develop into adults when their brains were dissected, processed, and analyzed for mistargeting as described above. Analysis was performed blinded to experimental manipulation. Control flies were the same genotype as experimental ones but reared exclusively at 18ºC.

### MARCM-based mosaic analysis

*hs-FLP* based MARCM analyses were performed by heat shocking 0–24h APF pupae for 1 hr at 37ºC. Each fly contained *AM29-GAL4, UAS-mCD8-GFP, UAS-mtdTomato, tubP-Gal80*, the desired *FRT* site, a mutant allele distal to the *FRT* site (or no allele for control), and in rescue experiments a *UAS-crb* rescue transgene (see Table S1 for complete genotypes). *ey-FLP* based MARCM analyses were not heat shocked but contained the same genomic components as *hs-FLP* MARCM flies. Brains were dissected, processed, and mounted as described above. MARCM phenotypes were analyzed blinded to genotype and a Fisher’s exact test was performed on MARCM phenotypes. P-values were adjusted using the Bonferroni correction.

### Transfection and immunostaining of *Drosophila* S2 cells

S2 cells were co-transfected with *Actin-GAL4* and either *pUASt-attB-Pvr*^*WT*^*-V5, pUASt-attB-Pvr*^*endo*.*mut*.^*-V5, pUASt-attB-HA-HRP*^*surf*^, *or pUASt-attB-HA-HRP*^*endo*^ using Effectine (Qiagen). After 48h, transfected cells were washed in 1X PBS, fixed in 4% PFA in PBS for 15 min, permeabilized in PBST, blocked in 5% normal donkey serum in PBST, and stained with: rabbit anti-HA (C29F4; Cell Signaling Technology) or rat anti-HA (3F10; Sigma Aldrich), mouse anti-Rab7 (Developmental Studies Hybridoma Bank), rabbit anti-Rab5 (Abcam); or mouse anti-V5 (ThermoFisher) or rabbit anti-V5 (ThermoFisher) at room temperature for 2 h, followed by three 15 min washes in PBST, and incubated in secondary antibody for 2 h at room temperature.

For antibody internalization assays, cells were transfected as previously described. After 48h, transfected cells were washed in 1X PBS and anti-HRP-594 was added to the media at 1:100. Cells were incubated at room temperature in media for 30, 60, 120, or 180 min, washed in PBS, fixed in 4% PFA in PBST, washed again three times in PBST for 15 min each, and imaged. The number of internalized HRP-594 puncta were quantified and normalized to the area of each cell.

### Gene ontology and STRING network analysis

Gene ontology (GO) analysis was performed using Flymine. We minimized GO redundancy using REVIGO ^86^. The STRING database was used to determine protein-protein interactions in Figure 4C, and nodes were connected based on experimental data. GO terms in chord and STRING plots were determined using Flymine. Specifically, we selected each term based on 1) breadth (we displayed broader terms); 2) experimental validation; and 3) neurodevelopmental relevance (we prioritized terms in this category). We displayed GO categories on these plots that had two or more proteins within them.

To generate the diagram in Figure 2E we compiled a list of proteins that have been shown to function in the endolysosomal system in mammalian cells ^8,91^ as well as in *Drosophila*. Since there has not been comprehensive profiling of the fly endolysosomal pathway we used Flybase (www.flybase.org) cellular component search tool to identify such proteins. For the mammalian proteins, we utilized the Flybase homologs search tool to identify the *Drosophila* orthologs of mammalian endolysosomal proteins.

### Quantification and statistical analysis

The statistical tests and numbers of independent replicates per experiment are indicated in the figures or figure legends.

### Quantification of co-localized puncta

Airyscan superresolution images (Carl Zeiss) were imported into Imaris10 (Oxford Instruments) and a mask was drawn to include the whole antennal lobe using the *Peb>FYVE-mCherry* channel. Volume renderings of puncta within each channel were subsequently made and the number of either Cont+ or Gli+ puncta co-localized with FYVE was computed. This value was divided by the total number of Cont+ or Gli+ puncta.

### Single-cell RNA-seq analysis

scRNA-seq data is from McLaughlin et al., 2021^3^ (ORNs) and Xie et al., 2021^5^ (PNs). For analysis in Figure 5, we averaged expression from the two developmental time points (24h APF and 48h APF) bracketing our profiling (36h APF) and from all ORN or PN subtypes, including clusters that have not yet been mapped to a specific ORN or PN subtype. A transcript was considered enriched if it was expressed in > 30% of cells at log_2_(CPM+1) ≥ 4. We note that a few transcripts that are only expressed in a few cell types or are very lowly expressed (Figure S7) may be false negatives in this analysis.

### Statistical analysis

Statistical tests for mutant phenotypic quantification are as follows. For categorical data we used a Fisher’s exact test followed by a Bonferroni correction to determine statistical significance. To determine statistical significance between continuous data sets, we used Mann-Whittney test for comparisons between two groups or a Krus-kal-Wallis test (followed by Dunn’s multiple comparisons test) for comparisons between multiple groups.

### Quantification of *Pvf3* expression in the antennal lobe

*Pvf3-GAL4* was crossed to *GH146-FLP; UAS>stop>mCD8-GFP* flies. Flies were reared at 25ºC and 42–48h APF pupae were dissected, labeled, and imaged according to the descriptions above. Glomeruli were identified based on stereotyped location and shape and fluorescence intensity in the brightest section of each glomerulus was measured. Background values were subtracted from intensity measurements and fluorescence intensity in each glomerulus was normalized to the average intensity in each antennal lobe. Fluorescence intensity of each glomerulus across all brains measured was averaged, and a heatmap was applied to each value. 3D volume renderings were created in Imaris10 (Oxford Instruments) and the heatmap values were transferred onto each glomerulus.

