## Supplemental Material for "Endocytome profiling uncovers cell-surface protein dynamics underlying neuronal connectivity"

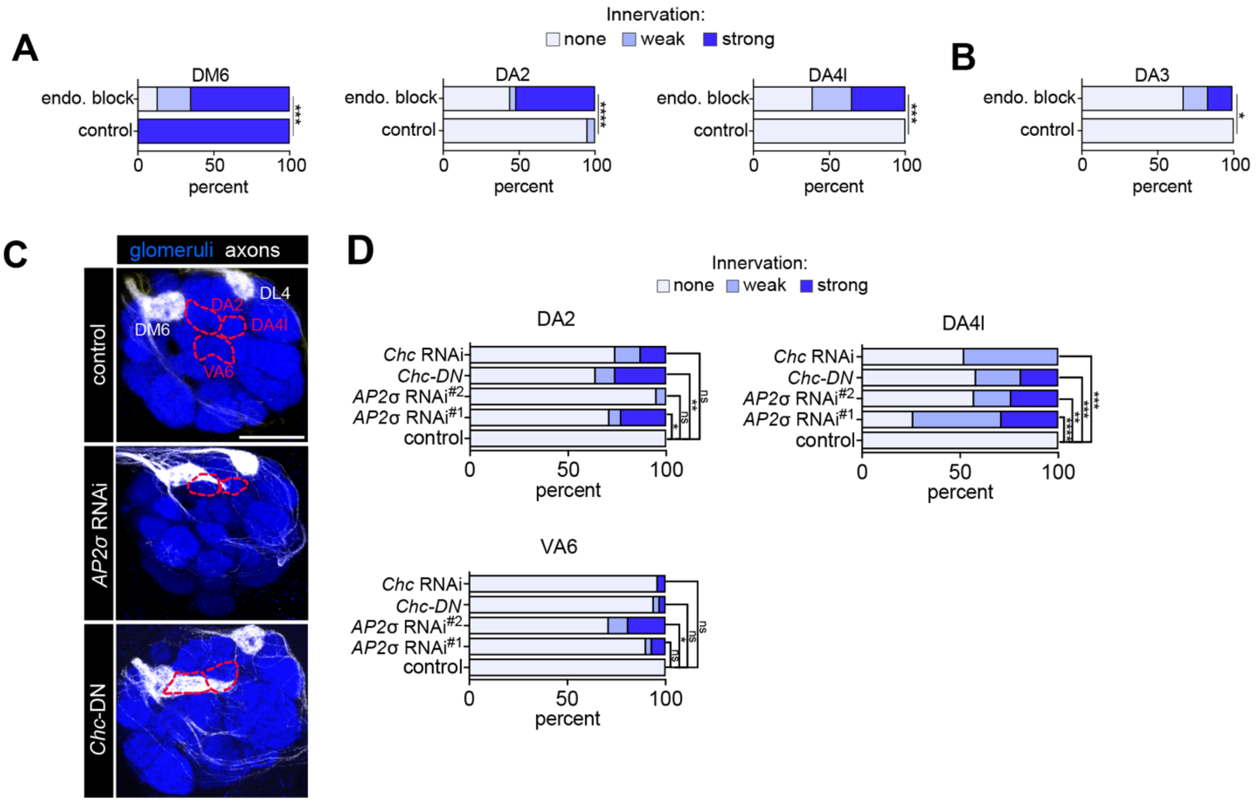

**Figure S1. Clathrin-mediated endocytosis is critical for axon targeting, related to Figure 1.**
(A, B) Quantification of axon targeting phenotype in DM6 (A) and DL4 (B) axons from Figure 1D, E. n = 23 (controls); n = 53 (endo. block).
(C) Images of axons innervating the DM6 and DL4 glomeruli (white, labeled with membrane tdTomato) of control (left) or those expressing UAS-RNAi against the σ-subunit of the AP2 complex (middle) or a dominant negative (DN) form of Clathrin heavy chain (Chc) (right) using the *AM29-GAL4* driver. Both AP2 and Chc mediate clathrin-dependent endocytosis of CSPs. Brains from late pupa were used due to adult lethality of some RNAi lines. Dotted outlines denote glomeruli where ectopic targeting is observed. Scale bar, 20 µm.
(D) Quantification of DM6 axon mistargeting phenotypes. n = 23 (control); n = 21 (*AP-2σ* RNAi[#1]); n = 31 (*AP-2σ* RNAi[#2]); n = 31 (Chc-DN); n = 23 (*Chc* RNAi).
Fisher's exact test was used to determine statistical significance.

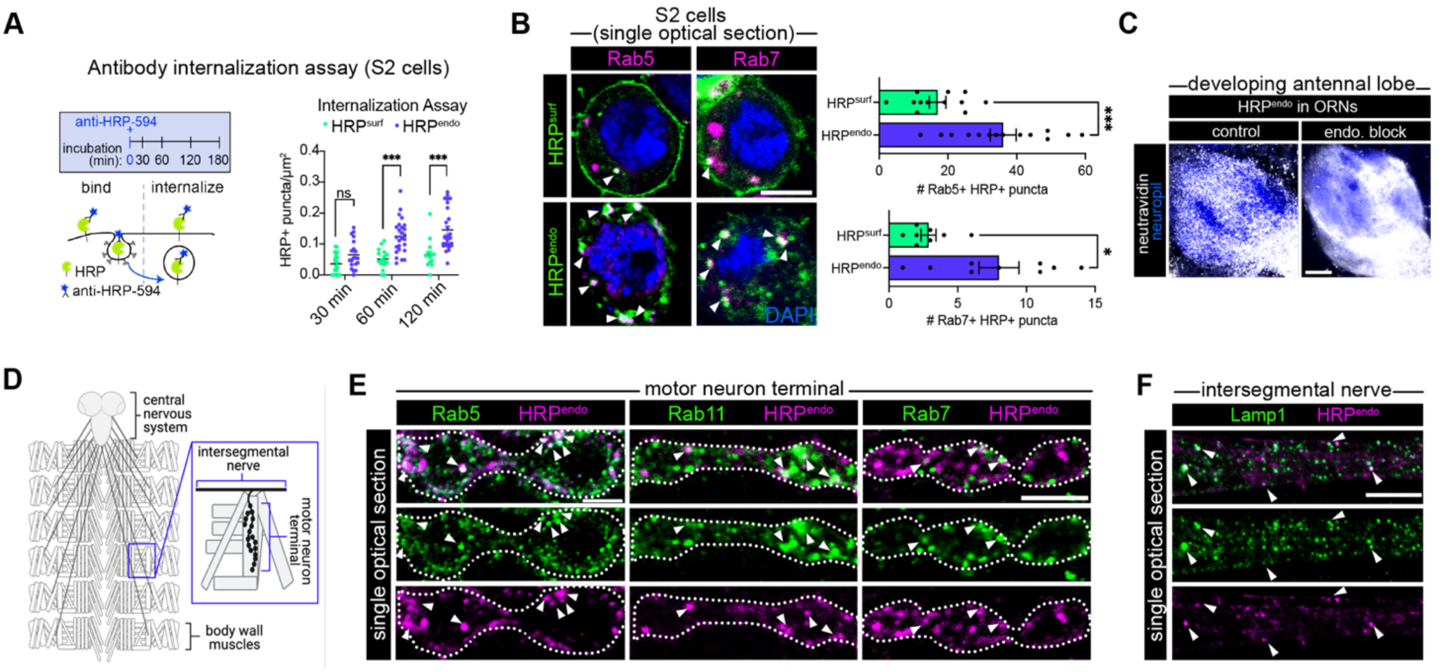

**Figure S2. Validation of endosome-targeted HRP, related to Figure 1.**

(A) Schematic of antibody internalization assay performed in S2 cells (left) and quantification of the number of internalized HRP-positive puncta per area (right). Note that some HRP$^{surf}$ is likely internalized at a low level due to its high surface expression.

(B) Images of S2 cells depicting co-labeling of indicated HRP and either Rab5 (early endosomes) or Rab7 (late endosomes) (left). Quantification of the number of HRP+ Rab+ puncta (right). Genotypes of all S2 cell experiments are either *UAS-HA-HRP$^{surf}$* or *UAS-HA-HRP$^{endo}$* driven by *Act-GAL4*.

(C) Images depicting the redistribution of neutravidin labeling from endosomes (left) to the surface (right) of ORN axons when endocytosis is blocked from 0–36h APF using *Peb>UAS-shi$^{ts}$*.

(D) Schematic of the third instar larval central nervous system and body wall muscles. Inset depicts the neuromuscular junction (NMJ). Motor neuron terminals were used to test colocalization because it is a large synapse of a single neuron, and colocalization can be performed *in vivo* with antibodies against endogenous proteins rather than overexpressed Rab/lysosomal proteins.

(E) Images of motor neuron terminals (NMJs) depicting colocalization between HRP$^{endo}$ and either Rab5, Rab11, or Rab7. Dotted outline denotes motor neuron membrane.

(F) Images of intersegmental nerves depicting colocalization between HRP$^{endo}$ and Lamp1.

Mann-Whitney test was used to determine statistical significance. Anti-HA was used to detect HRP transgenes unless otherwise noted. Scale bar, 5 μm (B, E), 10 μm (C, D). Arrowheads denote colocalization.

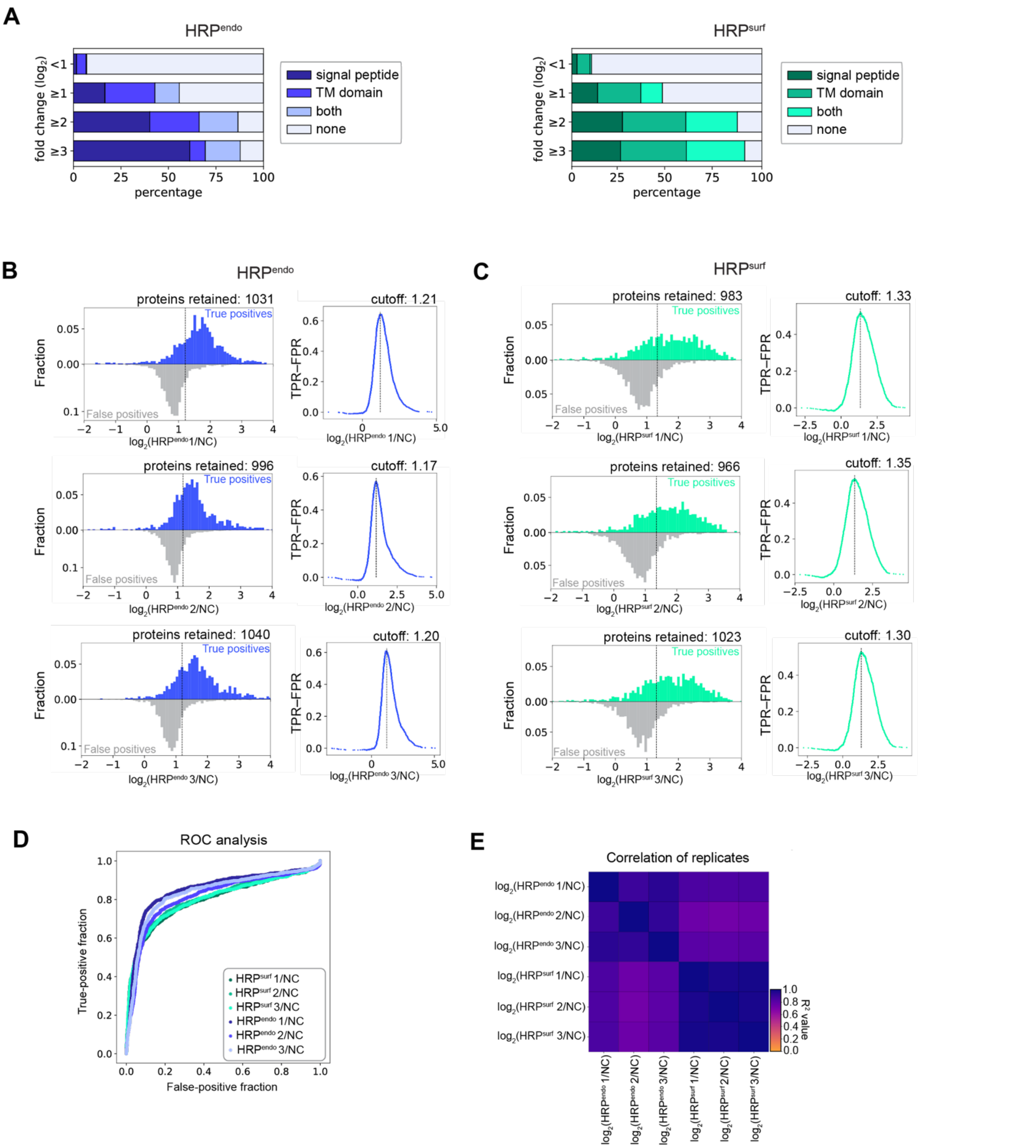

**Figure S3. Analysis of ORN endocytomes and surfaceomes, related to Figure 2.**
(A) Percentage of proteins labeled by $HRP^{endo}$ (left) or $HRP^{surf}$ (right) that contain a transmembrane (TM) domain or signal peptide (used to determine true-positive proteins). Fold change is compared to negative controls (NC).
(B, C) TMT-ratio cutoff in each biological replicate. Cutoffs were set to maximize true-positive rate – false-positive rate (TPR – FPR). True-positives are proteins with a signal peptide or TM domain and false-positives did not have these signatures and are annotated to the nucleus, mitochondria, or cytosol/cytoplasm by UniProt.

(D) Receiver operating characteristic (ROC) analysis for each biological replicate generated by plotting the proportion of true-positive proteins against the proportion of false-positive proteins rank ordered (from 0,0) by enrichment in each sample.
(E) Correlation matrix of biological replicates. Negative control (NC) data was averaged for this analysis.

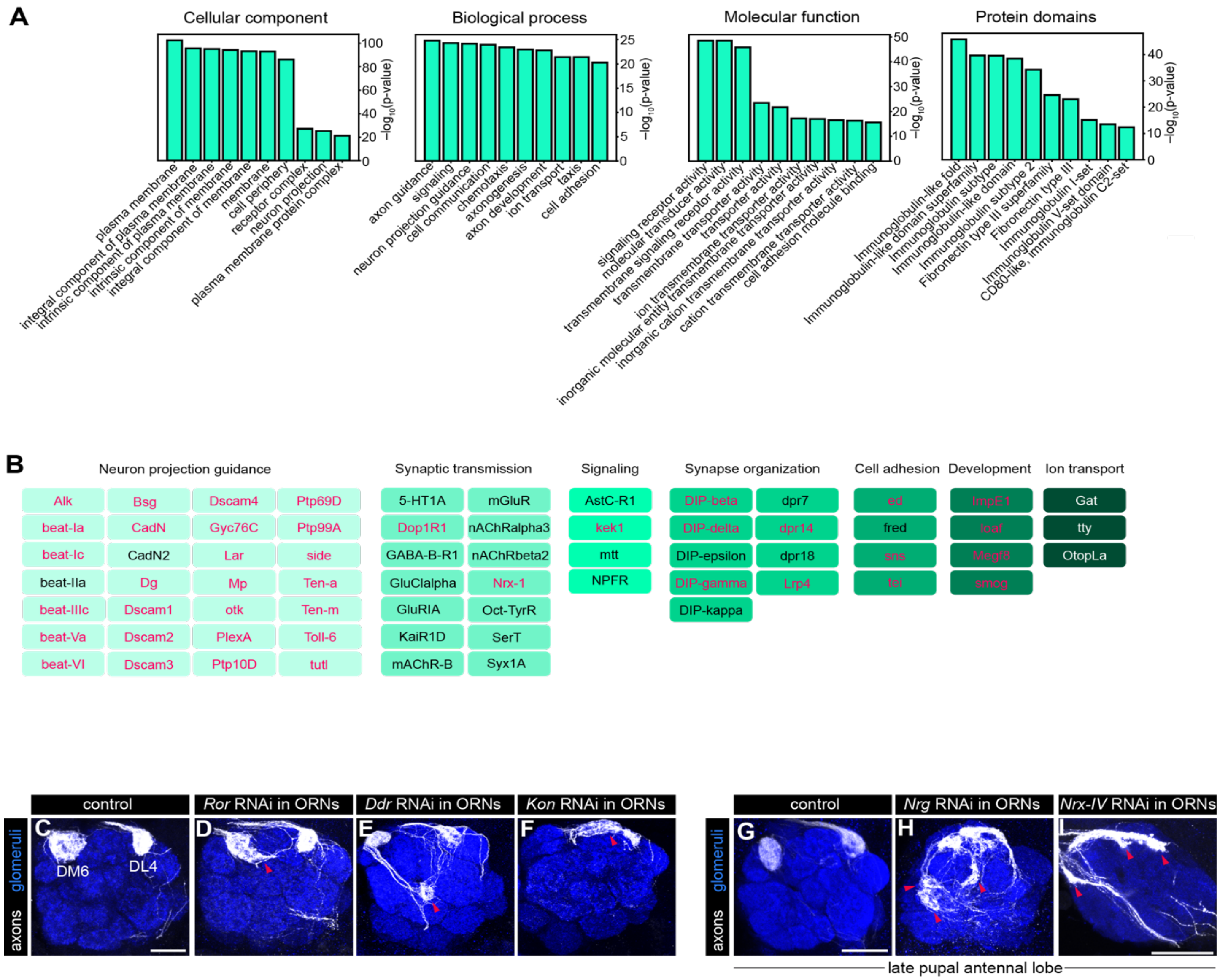

**Figure S4. Additional characterization of surface-enriched membrane proteins, related to Figure 4.**
(A) GO analysis of TM proteins present at the ORN axonal surface. Proteins with a $\log_2(\mathrm{HRP}^{\mathrm{surf}}/\mathrm{HRP}^{\mathrm{endo}}$ fold change$) \geq 0.1$ were used for this analysis.
(B) 66 of the top 100 CSPs enriched in surfaceomes and the biological process they are most associated with. For simplicity, we prioritized displaying GO categories that had more than 2 proteins annotated to them. The complete list of proteins, including unannotated ones, can be found in Table S2. Pink text indicates that these proteins have known neurodevelopmental functions according to Flymine GO evidence codes.
(C–I) Images of antennal lobes from the loss-of-function screen where membrane proteins enriched in ORN surfaceomes were knocked-down in all ORNs (via *peb-GAL4)* and axons projecting to the DM6 and DL4 glomerulus were labeled with *AM29-QF2* driven membrane-targeted tdTomato (*QUAS-mtdTomato*; white). Arrowheads indicate DM6-ORN axon mistargeting. Note for the two RNAi lines that caused lethality at the adult stage, we analyzed phenotypes in the late (~72–96h APF) pupal stages (K–M). Quantification of phenotypic penetrance is in Table S3.

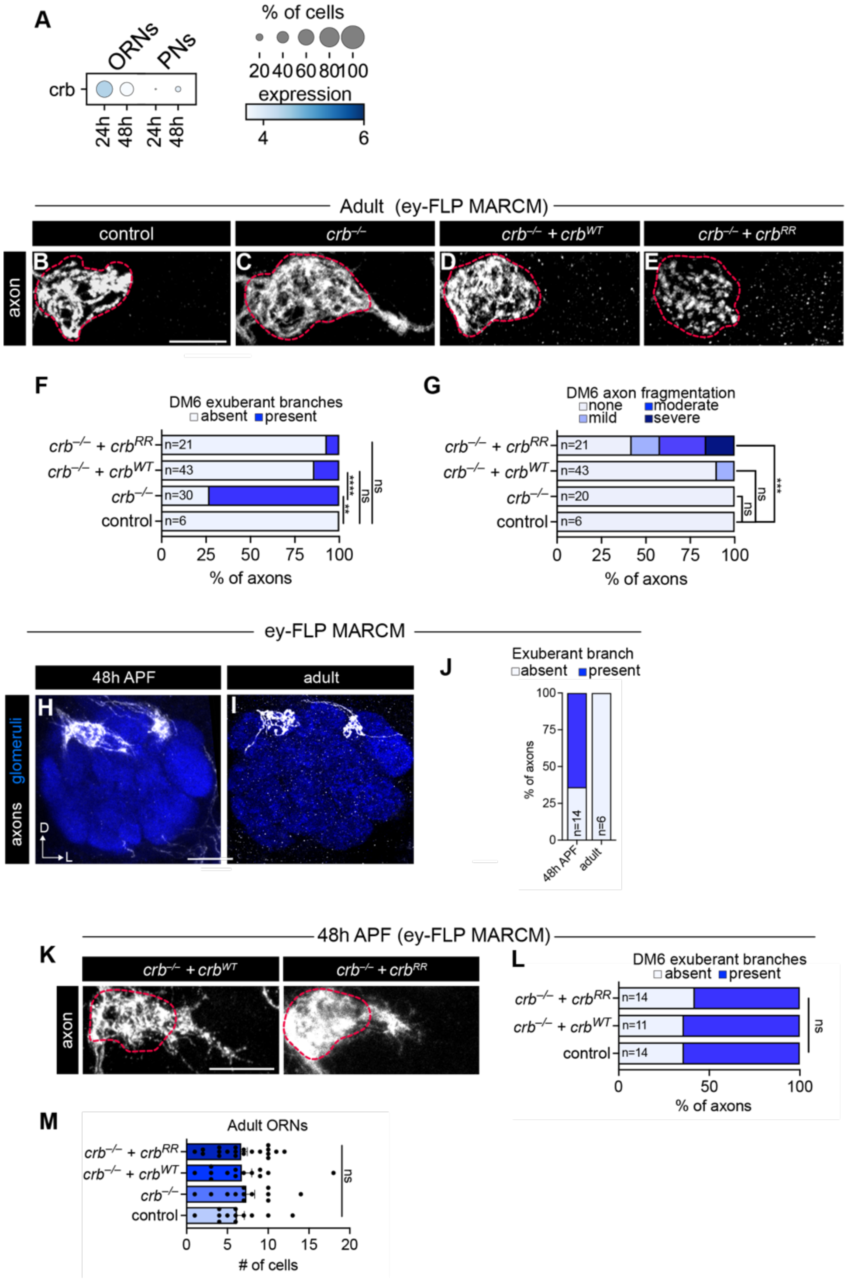

**Figure S5. Additional MARCM-based mosaic analysis of *crb* mutant backgrounds, related to Figure 5.**
(A) Dotplot depicting expression of *crb*. Expression unit is log$_2$(CPM+1).
(B–E) Images of mosaic analysis using *ey-FLP* MARCM of DM6 axon phenotypes in indicated genotypes.
(F, G) Quantification of the percentage of DM6 axons that still have the exuberant branches present at the adult stage (E) or have axon fragmentation (F) in indicated genotypes.
(H, I) Images of *ey-FLP* DM6 axon clones during development (48h APF) and in the adult stage.
(J) Quantification of the number of antennal lobes where DM6 axons have exuberant branches at indicated time points.
(K) Images depicting exploring branches in DM6 axons of *ey-FLP* MARCM *crb* LOF clones that express either *crb* rescue construct, *UAS-GFP-*$crb^{WT}$ (left) or *UAS-GFP-*$crb^{RR}$ (right).
(L) Quantification of the number of antennal lobes where DM6 axons have exuberant branches in indicated genotypes.
(M) Quantification of the number of cell bodies present in the adult antenna of *ey-FLP* MARCM clones in indicated genotypes. Dashed lines denote DM6 glomerular boundary. Scale bar, 20 µm. Fisher's exact test (F, G, L) and Kruskal-Wallis test (M) was used to determine statistical significance.

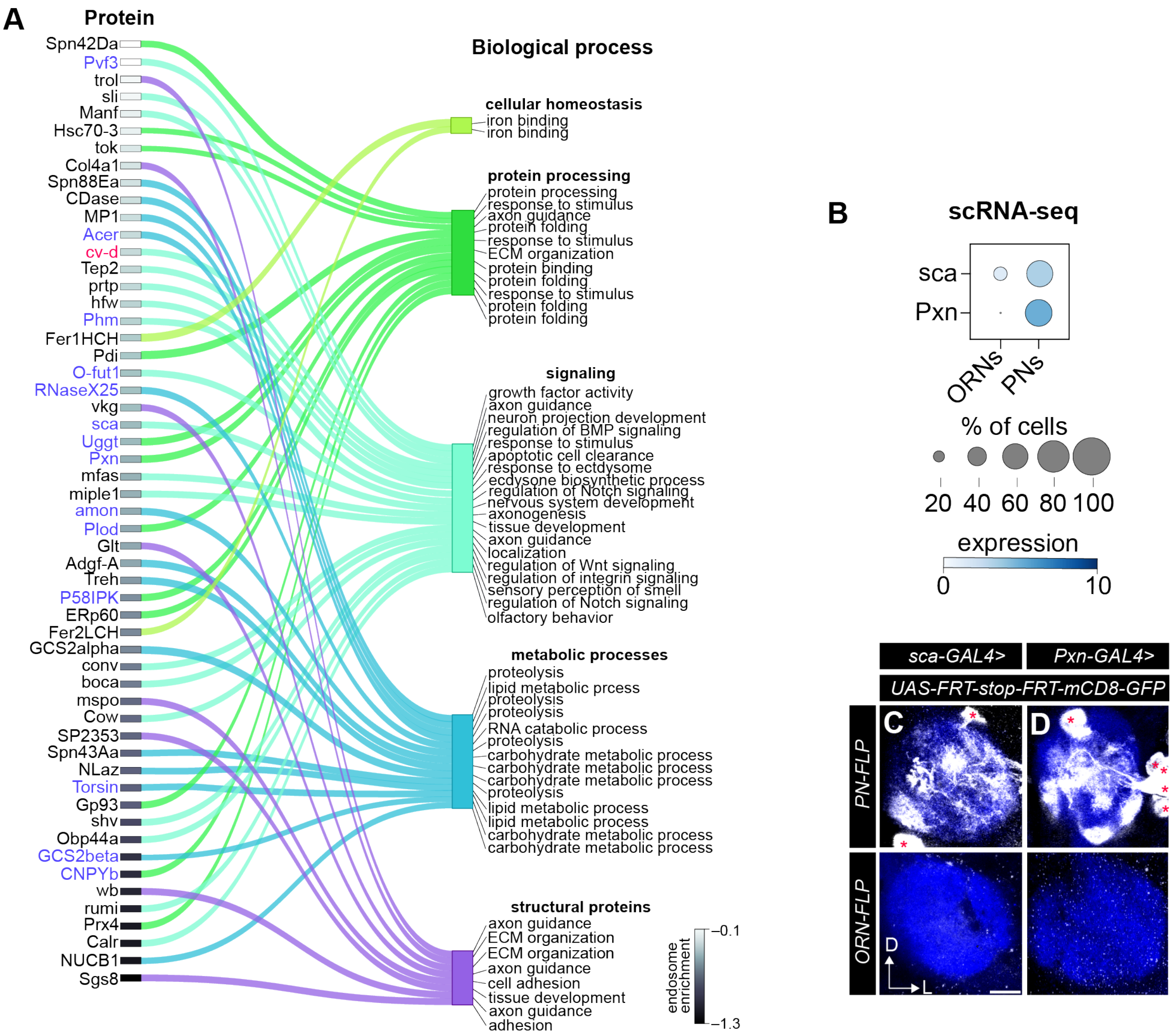

**Figure S6. Depiction and validation of cell-type specific expression of endosome-enriched secreted proteins, Related to Figure 6.**

(A) Chord plot depicting endosome-enriched secreted proteins (left) and the biological process they are associated with (right). Proteins are rank ordered based on endosomal enrichment (highest at the bottom) with the grayscale in the rectangles to the right of protein names indicating the level of enrichment ($\log_2$[HRP$^{surf}$/HRP$^{endo}$] fold change). See STAR Methods for details on GO categories. Detailed protein information can be found in Table S1. Transcripts that are highly expressed in PNs and ORNs (by scRNA-seq) are in blue and pink, respectively.

(B) Dotplot depicting scRNA-seq data showing that *sca* and *Pxn* are preferentially expressed in PNs.

(C, D) Images of ~36h APF antennal lobes depicting GAL4 expression used to validate cell-type-specific expression found in (A). Asterisks denote PN cell bodies. Scale bar, 10 µm.

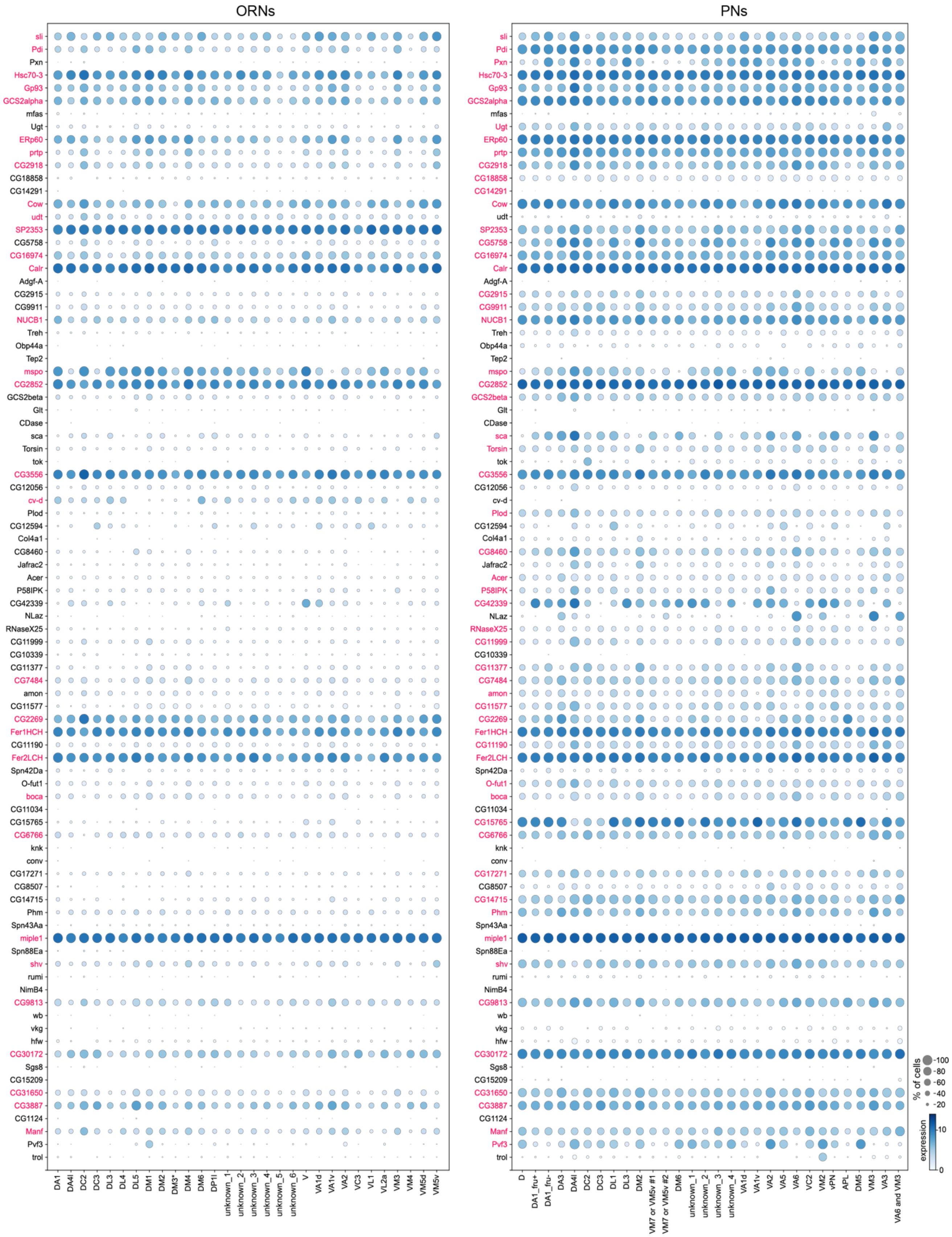

**Figure S7. Single-cell RNA-sequencing analysis of genes encoding endocytome-enriched secreted proteins, related to Figure 6.** Dotplots depicting expression of transcripts encoding endocytome-enriched secreted proteins in ORNs (left) and PNs (right). Expression was averaged across two developmental timepoints (24h APF and 48h APF). Expression is in

$\log_2$(CPM + 1). Transcripts in pink met expression threshold ($\log_2$[CPM+1] ≥ 4 in ≥ 30% of cells) and were retained for further analysis. 'unknown_#' denotes a cluster where the glomerular target of an ORN or PN subtype is unknown.

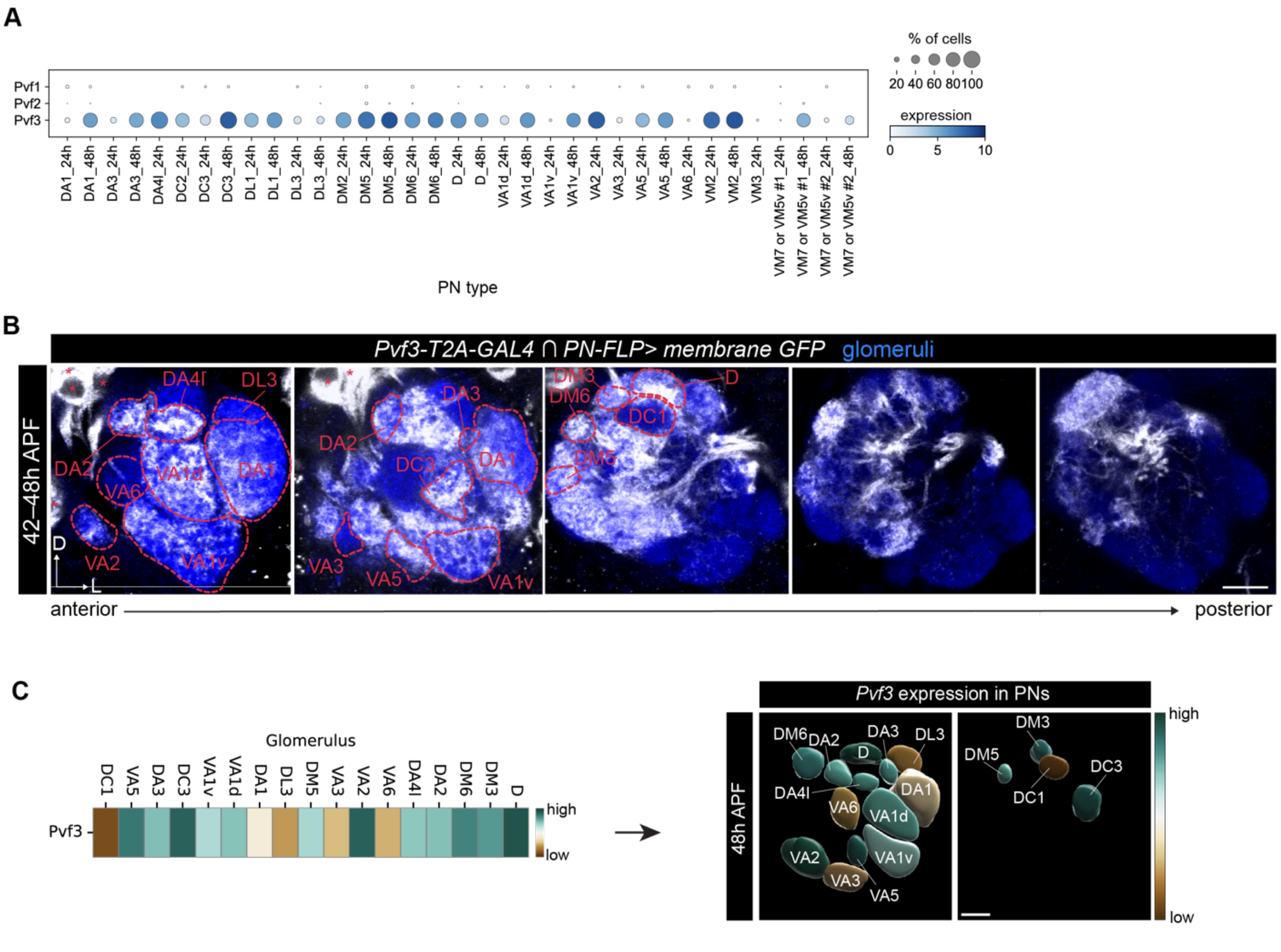

**Figure S8. *Pvf3* expression analyses, related to Figure 7.**
(A) Dotplot depicting expression of mRNAs (*Pvf1–3*) that encode Pvr's known ligands in 48–54h APF PNs. Expression unit is $\log_2$(CPM+1).
(B) Images along the anteroposterior axis of the developing antennal lobe of *Pvf3-T2A-GAL4* expression in PNs. Fluorescence intensity of encircled glomeruli was used to generate plots in Figure 4C. Asterisks denote PN cell bodies. Slight differences in transcript expression between (A) and (B) likely reflect differences in the stages where expression was analyzed (scRNA-seq is from 48-54h APF and fluorescence is from 42-48h APF). Scale bar, 10 µm.
(C) Heatmaps depicting averaged fluorescence intensity in each glomerulus of GAL4 lines in B. Fluorescence intensity in each glomerulus was normalized to the average intensity of all analyzed glomeruli. n ≥ 5 brains were analyzed for each GAL4 line. Heatmap colors were directly used to generate the overlay colors in Figure 7G. Expression is in arbitrary units (left). Volume rendering of a subset of surface glomeruli in the antennal lobe depicting *Pvf3* expression (right).

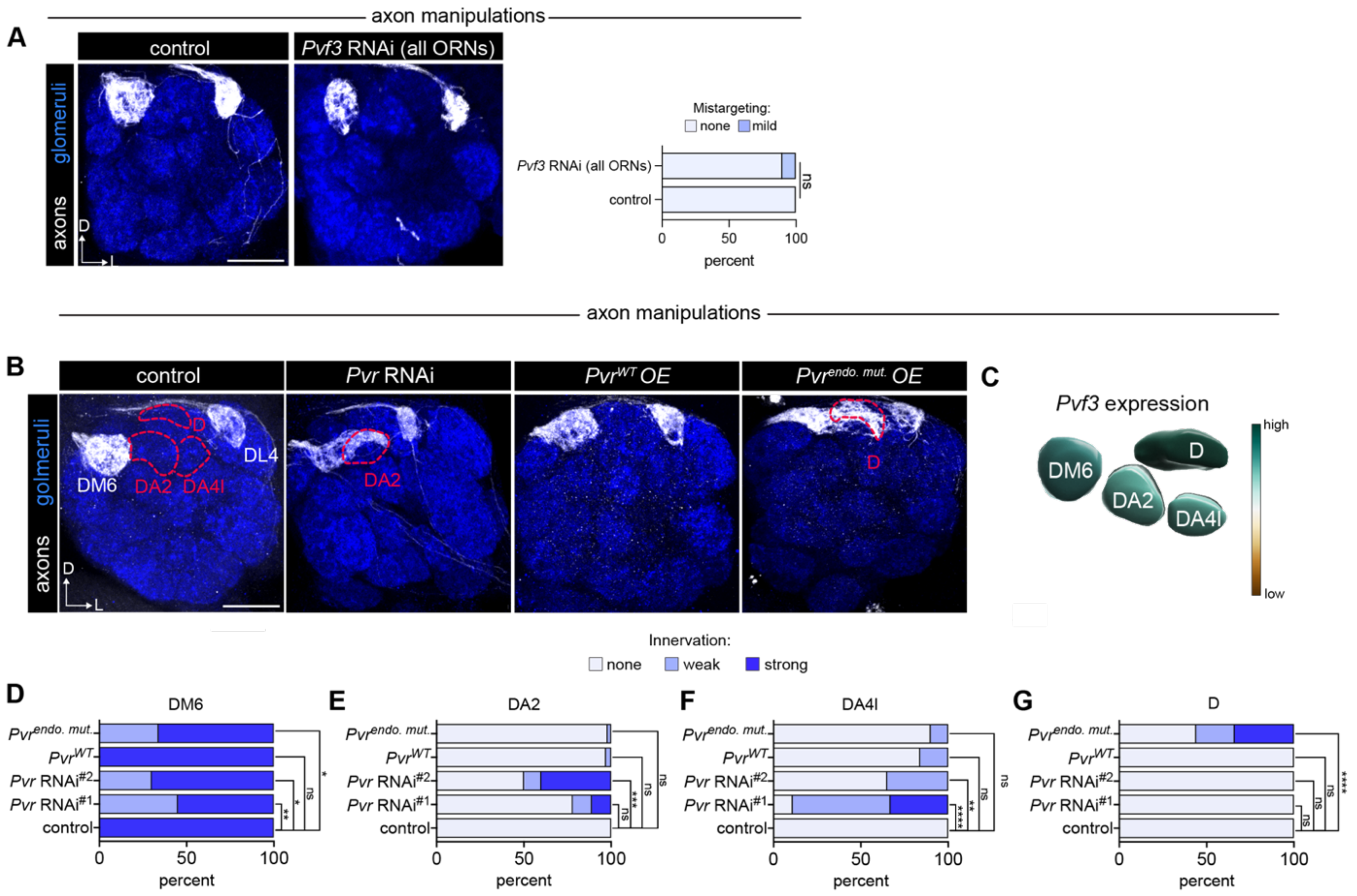

**Figure S9. Extended analyses of axon targeting in *Pvf3* and *Pvr* loss- and gain-of-function manipulations in DM6 ORNs, related to Figure 7.**
(A) Representative images of DM6-ORN and DL4-ORN axons of controls or when *Pvf3* is knocked down in all ORNs illustrating that there is no phenotype when the ligand is decreased in ORNs (left) and quantification of mistargeting in indicated genotypes (right). n = 19 (controls); n = 30 (*Pvf3* RNAi*)*.
(B) Representative images of DM6-ORN axons in indicated genotypes. Encircled glomeruli denote where ectopic targeting occurs when *Pvr* is manipulated.
(C) Volume rendering depicting *Pvf3* expression in glomeruli analyzed in subsequent panels.
(D–G) Quantification of DM6-ORN axon mistargeting phenotypes. Fisher's exact test was used to determine statistical significance. n = 23 (controls); n = 18 (*Pvr RNAi*$^{\#1}$); n = 20 (*Pvr RNAi*$^{\#2}$); n = 31 (*Pvr*$^{WT}$ OE); n = 41 (*Pvr*$^{endo.mut.}$ OE). Scale bar, 20 µm.

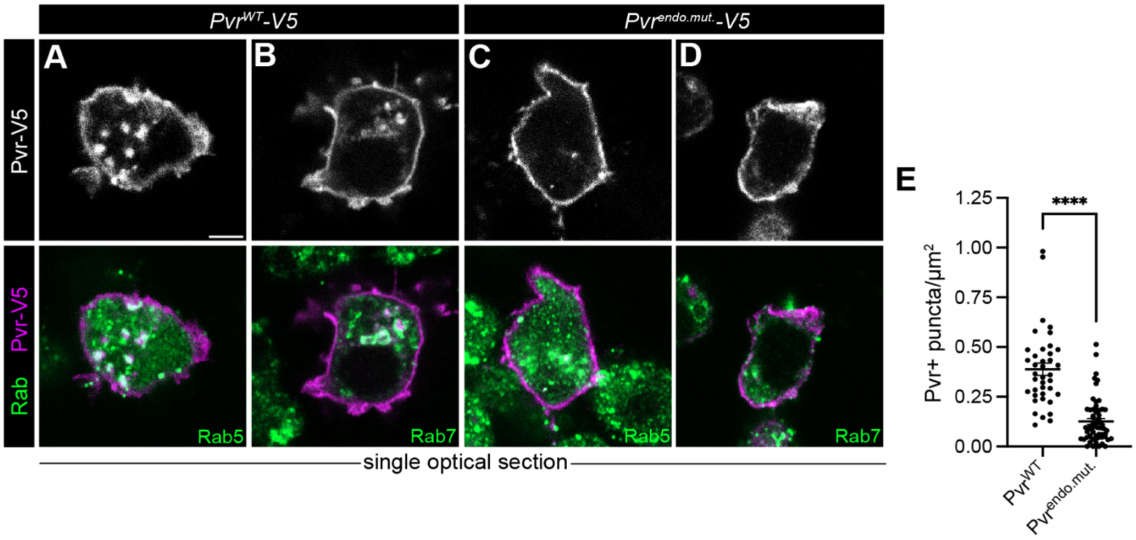

**Figure S10. Extended analyses of Pvr overexpression transgenes, related to Figure 7.**
(A–D) Images of S2 cells depicting co-labeling of indicated Pvr-V5 transgene and either Rab5 (A, C) or Rab7 (B, D).
(E) Quantification of the number of intracellular V5+ puncta per area. Number of intracellular puncta was normalized to the cell area to account for differences in cell size. Genotypes of experiments are either *UAS-Pvr$^{WT}$-V5* or *UAS-Pvr$^{endo.mut.}$-V5* driven by *Act-GAL4*. Scale bar is 5 µm. Kruskal-Wallis test was used to determine statistical significance. Scale bar is 2 µm.

| Figure | Genotype | Notes |
|---|---|---|
| **Figure 1** | | |
| **D,E** | *AM29-GAL4, UAS-S-mtdTomato-3xHA/+; 20XUAS-IVS-shits1-p10*_BDSC 66599 | control = 18ºC; endo. Block = 24h @ 30ºC |
| **Figure 2** | | |
| **C, D** | *Peb-GAL4/+; UAS-HRP^endo/+* | |
| **E, F** | *Peb-GAL4/+; UAS-HRP^surf/+* | |
| **H** | *Peb-GAL4/+; UAS-HRP^endo/+* (endosome) | |
| | *Peb-GAL4/+; UAS-HRP^surf/+* (surface) | |
| | *w1118* (no enzyme control) | |
| | *Peb-GAL4/+; UAS-HRP^endo/+* (no hydrogen peroxide control) | |
| **Figure 3** | | |
| **D** | *Peb-GAL4/+; UAS-mCherry-2xFYVE/+* | |
| | *Peb-GAL4/+; UAS-mCherry-2xFYVE/Gli-YFP* | |
| **Figure 4** | | |
| **E** | *Peb-GAL4, UAS-Dcr2/+;AM29-QF2, QUAS-mtdTomato-3xHA/+; UAS-luc*_BDSC 35788 | |
| **F** | *Peb-GAL4, UAS-Dcr2/+;AM29-QF2, QUAS-mtdTomato-3xHA/+; UAS-bark RNAi*_VDRC 52608/+ | |
| **G** | *Peb-GAL4, UAS-Dcr2/+;AM29-QF2, QUAS-mtdTomato-3xHA/+; UAS-crb RNAi*_BDSC 27697/+ | |
| **H** | *Peb-GAL4, UAS-Dcr2/+;AM29-QF2, QUAS-mtdTomato-3xHA/UAS-Gli RNAi*_BDSC 38284 | |
| **I** | *Peb-GAL4, UAS-Dcr2/+;AM29-QF2, QUAS-mtdTomato-3xHA/UAS-Tsf2 RNAi*_BDSC 65903 | |
| **J** | *Peb-GAL4, UAS-Dcr2/+;AM29-QF2, QUAS-mtdTomato-3xHA/UAS-pck RNAi*_BDSC 38203 | |
| **K** | *Peb-GAL4, UAS-Dcr2/+;AM29-QF2, QUAS-mtdTomato-3xHA/UAS-sinu RNAi*_VDRC 44928 | |
| **Table S3** | *Peb-GAL4, UAS-Dcr2/+; AM29-QF2, QUAS-mtdTomato-3xHA/+ x* UAS-RNAi lines | Line numbers and stock centers are in table |
| **Figure 5** | | |
| **B** | *Peb-GAL4; UAS-mCherry-2xFYVE/+; Crb-C-GFP/+* | |
| **C** | *hs-FLP, UAS-mCD8-GFP/+;AM29-GAL4,UAS-mtdTomato-3xHA/+; FRT82b, TubP-GAL80/FRT82b* | |
| **D** | *hs-FLP, UAS-mCD8-GFP/+;AM29-GAL4,UAS-mtdTomato-3xHA/+; FRT82b, TubP-GAL80/FRT82b, crb^11A22* | |
| **E** | *hs-FLP, UAS-mCD8-GFP/+;AM29-GAL4,UAS-mtdTomato-3xHA/+; FRT82b, TubP-GAL80/UAS-GFP-crb^WT,FRT82b, crb^11A22* | |
| **F** | *hs-FLP, UAS-mCD8-GFP/+;AM29-GAL4,UAS-mtdTomato-3xHA/+; FRT82b, TubP-GAL80/UAS-GFP-crb^RR,FRT82b, crb^11A22* | |
| **I** | *AM29-GAL4, UAS-mtdTomato-3xHA/+; UAS-GFP-crb^WT/+* | |
| **J** | *AM29-GAL4; UAS-mChery-2xFYVE/UAS-GFP-crb^WT* | |
| **K** | *AM29-GAL4; UAS-mChery-2xFYVE/UAS-GFP-crb^RR* | |
| **Figure 6** | | |
| **G** | *AM29-QF2, QUAS-mtdTomato-3xHA/ UAS-luc; VT033006-Gal4/+* | BDSC_35788 |
| **H** | *AM29-QF2, QUAS-mtdTomato-3xHA/; VT033006-Gal4/UAS-shv-RNAi*_VDRC 22997 | BDSC_54797 (#1)<br>VDRC_22997 (#2) |
| **I** | *AM29-QF2, QUAS-mtdTomato-3xHA/ UAS-Pvf3-RNAi*_VDRC 105008; *VT033006-Gal4/+* | VDRC_105008 (#1)<br>BDSC_38962 (#2) |
| **Figure 7** | | |
| **B, C** | *UAS-dcr2, UAS-mCD8-GFP/+; Mz19-GAL4, UAS-mCD8-GFP, Or88a-mtdTomato/+; Or47b-rCD2/UAS-*luc_BDSC 35788 | |

| | | |
|---|---|---|
| **D** | *UAS-dcr2, UAS-mCD8-GFP/+; Mz19-GAL4, UAS-mCD8-GFP, Or88a-mtdTomato/UAS-Pvf3-RNAi_*VDRC 105008*; Or47b-rCD2/+* | VDRC_105008 (#1)<br>BDSC_38962 (#2) |
| **E** | *UAS-dcr2, UAS-mCD8-GFP/+; Mz19-GAL4, UAS-mCD8-GFP, Or88a-mtdTomato/+; Or47b-rCD2/UAS_Pvf3-myc* | |
| **H** | *UAS-dcr2, UAS-mCD8-GFP/+; R20D10-QF2, QUAS-mtdTomato-3xHA/+; R78H05-p65AD, R31F09-GAL4DBD/UAS-luc_*BDSC 35788 | Did not display labeling for R20D10>mtdTomato in I–L |
| **I** | *UAS-dcr2, UAS-mCD8-GFP/+; R20D10-QF2, QUAS-mtdTomato-3xHA/UAS-Pvr RNAi_*VDRC 977 | VDRC_977 (#1); BDSC_37520 (#2) |
| **J** | *UAS-dcr2, UAS-mCD8-GFP/+; R20D10-QF2, QUAS-mtdTomato-3xHA/+; R78H05-p65AD, R31F09-GAL4DBD/UAS-Pvr^WT-V5* | |
| **K** | *UAS-dcr2, UAS-mCD8-GFP/+; R20D10-QF2, QUAS-mtdTomato-3xHA/+; R78H05-p65AD, R31F09-GAL4DBD/UAS-Pvr^endo.mut.-V5* | |
| **Q, R** | *AM29-GAL4, UAS-mtdTomato-3xHA/+; UAS-Pvr^WT-V5/+* | |
| **S, T** | *AM29-GAL4, UAS-mCherry-2xFYVE/+; UAS-Pvr^WT-V5/+* | |
| **Figure S1** | | |
| **A,B** | *AM29-GAL4, UAS-mtdTomato-3xHA/+; UAS-luc_*BDSC 35788/+ (control) | |
| | *AM29-GAL4, UAS-mtdTomato-3xHA/+; UAS-AP-2sigma_*BSDC 27322/+ | |
| | *UAS-AP-2sigma_*VDRC 110725/+*; AM29-GAL4, UAS-mtdTomato-3xHA/+* | |
| | *AM29-GAL4, UAS-mtdTomato-3xHA/+; UAS-Chc-DN_*BDSC 26874/+ | |
| | *AM29-GAL4, UAS-mtdTomato-3xHA/+; UAS-Chc RNAi_*BSDC 34742/+ | |
| **Figure S2** | | |
| **C** | *Peb-GAL4, ey-FLP/+; UAS-FRT-stop-FRT-shi(ts1)/UAS-HRP^endo; UAS-FRT-stop-FRT-shi(ts1)/+* | control = 18ºC; endo. block = 30ºC |
| **E, F** | *OK6-GAL4/UAS-HRP^endo* | |
| **Figure S4** | | |
| **C** | *Peb-GAL4, UAS-Dcr2/+;AM29-QF2, QUAS-mtdTomato-3xHA/+; UAS-luc_*BDSC 35788 | |
| **D** | *Peb-GAL4, UAS-Dcr2/+;AM29-QF2, QUAS-mtdTomato-3xHA/UAS-Ror RNAi_*BDSC 62868 | |
| **E** | *Peb-GAL4, UAS-Dcr2/+;AM29-QF2, QUAS-mtdTomato-3xHA/+; UAS-Ddr RNAi _*VDRC 51719/+ | |
| **F** | *Peb-GAL4, UAS-Dcr2/+;AM29-QF2, QUAS-mtdTomato-3xHA/+; UAS-Kon RNAi_*BDSC 31584/+ | |
| **G** | *Peb-GAL4, UAS-Dcr2/+;AM29-QF2, QUAS-mtdTomato-3xHA/+; UAS-luc_*BDSC 35788 | |
| **H** | *Peb-GAL4, UAS-Dcr2/+;AM29-QF2, QUAS-mtdTomato-3xHA/UAS-Nrg RNAi_*BDSC 38215 | |
| **I** | *Peb-GAL4, UAS-Dcr2/+; AM29-QF2, QUAS-mtdTomato-3xHA/+; UAS-Nrx-IV RNAi_*BDSC 32424 | |
| **Figure S5** | | |
| **B** | *ey-FLP, UAS-mCD8-GFP/+; AM29-GAL4, UAS-mtdTomato-3xHA/+; FRT82b, TubP-GAL80/FRT82b* | |
| **C** | *ey-FLP, UAS-mCD8-GFP/+; AM29-GAL4, UAS-mtdTomato-3xHA/+; FRT82b, TubP-GAL80/FRT82b, crb^11A22* | |
| **D** | *ey-FLP, UAS-mCD8-GFP/+; AM29-GAL4, UAS-mtdTomato-3xHA/+; FRT82b, TubP-GAL80/UAS-GFP-crb^WT,FRT82b, crb^11A22* | |
| **E** | *ey-FLP, UAS-mCD8-GFP/+; AM29-GAL4, UAS-mtdTomato-3xHA/+; FRT82b, TubP-GAL80/UAS-GFP-crb^RR,FRT82b, crb^11A22* | |
| **H, I** | *ey-FLP, UAS-mCD8-GFP/+; AM29-GAL4, UAS-mtdTomato-3xHA/+; FRT82b, TubP-GAL80/FRT82b* | |
| **K** | *ey-FLP, UAS-mCD8-GFP/+; AM29-GAL4, UAS-mtdTomato-3xHA/+; FRT82b, TubP-GAL80/UAS-GFP-crb^WT,FRT82b, crb^11A22* (left) | |
| | *ey-FLP, UAS-mCD8-GFP/+; AM29-GAL4, UAS-mtdTomato-3xHA/+; FRT82b, TubP-GAL80/UAS-GFP-crb^RR,FRT82b, crb^11A22* (right) | |
| **Figure S6** | | |
| **C** | *GH146-FLP, UAS-FRT-stop-FRT-mCD8-GFP/sca-GAL4_*BDSC 6479 (top) | |
| | *ey-FLP3.5/+; UAS-FRT-stop-FRT-mCD8-GFP/sca-GAL4_*BDSC 6479 (bottom) | |

| | |
|---|---|
| **D** | *GH146-FLP, UAS-FRT-stop-FRT-mCD8-GFP/+; Pxn-GAL4/+*_BDSC 66850 (top) |
| | *ey-FLP3.5/+; UAS-FRT-stop-FRT-mCD8-GFP/+; Pxn-GAL4/+*_BDSC 66850 (bottom) |
| **Figure S8** | |
| **B** | *GH146-FLP, UAS-FRT-stop-FRT-mCD8-GFP/Pvf3-T2A-GAL4* |
| **Figure S9** | |
| **A** | *Peb-GAL4, UAS-Dcr2/+; AM29-QF2, QUAS-mtdTomato-3xHA/+; UAS-luc*_BDSC 35788 (left) |
| | *Peb-GAL4, UAS-Dcr2/+; AM29-QF2, QUAS-mtdTomato-3xHA/UAS-Pvf3-RNAi*_VDRC 105008 (right) |
| **B–G** | *AM29-GAL4, UAS-mtdTomato-3xHA/+; UAS-luc*_BDSC 35788/+ (control) |
| | *AM29-GAL4, UAS-mtdTomato-3xHA/+; UAS-Pvr RNAi*_VDRC 977 (#1) |
| | *AM29-GAL4, UAS-mtdTomato-3xHA/ UAS-Pvr RNAi*_BDSC 37520 (#2) |
| | *AM29-GAL4, UAS-mtdTomato-3xHA/UAS-Pvr^WT-V5* |
| | *AM29-GAL4, UAS-mtdTomato-3xHA/UAS-Pvr^endo.mut.-V5* |

**Table S1.** Complete list of genotypes used in each figure.

**Table S2**. Processed proteomics data, related to Figure 2, S3, 5, S4. See Excel spreadsheet.

| Protein | Enrichment | Human ortholog(s) | DM6-ORN axon targeting deficits caused by RNAi knockdown | % Phenotypic penetrance (n) | RNAi lines |
|---|---|---|---|---|---|
| **bark** | endosome | – | axon trajectory error | 27 (22); 56 (16) | BDSC 67014; VDRC 52608 |
| **crb** | endosome | CRB1, CRB2 | local mistargeting | 20 (30); 74 (23) | BDSC 34999; BDSC 27697 |
| **Gli** | endosome | NLGN1–3 | local mistargeting | 73 (33); 68 (31) | BDSC 58115; BDSC 38284 |
| **pck** | endosome | Claudin | long-range mistargeting | 79 (24); 18 (16) | BDSC 38203; VDRC50306 |
| **sinu** | endosome | Claudin | long-range mistargeting | 60 (30); 82 (17) | BDSC 38200; VDRC 44928 |
| **Tsf2** | endosome | MELTF | local mistargeting | 61 (28) | BDSC 65903 |
| **Ddr** | surface | DDR1, DDR2 | local and long-range mistargeting | 31 (39); 81 (16) | VDRC 51719; BDSC 55906 |
| **Kon** | surface | CSPG4 | local mistargeting | 64 (36); 52 (27) | BDSC 31583; BDSC 80477 |
| **Nrg** | surface | NRCAM, L1CAM, CHL1, NFASC | local and long-range mistargeting; DM6 & DL4 merge; loss of bilateral innervation* | 98 (40) | BDSC 38215 |
| **Nrx-IV** | surface | CNTNAP1–3 | loss of glomerular and terminal morphology; DM6 & DL4 terminals merge | 79 (24); 77 (26) | BDSC 32424; VDRC 108128 |
| **Ror** | surface | ROR1, ROR2 | local mistargeting | 73 (46); 41 (37) | BDSC 62868; VDRC 29930 |
| **Drl-2** | surface | RYK | none | – | BDSC 55983; VDRC 40484 |

**Table S3, related to Figure 4 and S4.** A subset of endosome- and surface-enriched proteins were knocked-down in all ORNs (via *peb-GAL4)* and targeting to the DM6 glomerulus (using *AM29QF2, QUAS-mtdTomato*) was evaluated. Human orthologs were identified using Flybase Homologs search tool. The most predominant DM6 axon phenotype(s) is described with the penetrance of each RNAi line listed and the number of antennal lobes examined in parentheses. Note that *peb-GAL4* is expressed in tissues outside of the nervous system and when used to drive certain RNAi lines caused lethality, thus two proteins have a single RNAi line used for analysis. *Indicates that the loss of bilateral innervation caused by Nrg RNAi was previously described[94].
